# The Modular µSiM: a Mass Produced, Rapidly Assembled, and Reconfigurable Platform for the Study of Barrier Tissue Models *In Vitro*

**DOI:** 10.1101/2022.03.28.486095

**Authors:** Molly C. McCloskey, Pelin Kasap, S. Danial Ahmad, Shiuan-Haur Su, Kaihua Chen, Mehran Mansouri, Natalie Ramesh, Hideaki Nishihara, Yury Belyaev, Vinay V. Abhyankar, Stefano Begolo, Benjamin H. Singer, Kevin F. Webb, Katsuo Kurabayashi, Jonathan Flax, Richard E. Waugh, Britta Engelhardt, James L. McGrath

## Abstract

Advanced *in vitro* tissue chip models can reduce and replace animal experimentation and may eventually support ‘on-chip’ clinical trials. To realize this potential, however, tissue chip platforms must be both mass-produced and reconfigurable to allow for customized design. To address these unmet needs, we introduce an extension of our µSiM (**micro**device featuring a **si**licon-nitride **m**embrane) platform. The *modular* µSiM (m-µSiM) uses mass-produced components to enable rapid assembly and reconfiguration by laboratories without knowledge of microfabrication. We demonstrate the utility of the m-µSiM by establishing an hiPSC-derived blood-brain barrier (BBB) in bioengineering and non-engineering, brain barriers focused laboratories. We develop and validate *in situ* and sampling-based assays of small molecule diffusion as a measure of barrier function. BBB properties show excellent interlaboratory agreement and match expectations from literature, validating the m-µSiM as a platform for barrier models and demonstrating successful dissemination of components and protocols. We then demonstrate the ability to quickly reconfigure the m-µSiM for co-culture and immune cell transmigration studies through addition of accessories and/or quick exchange of components. Because the development of modified components and accessories is easily achieved, custom designs of the m-µSiM should be accessible to any laboratory desiring a barrier-style tissue chip platform.

## 1. Introduction

Negative pressures on the use of animal models in medical research have emerged from animal welfare advocacy and Russell’s 3R’s: *reduce*, *refine*, and *replace,* guiding the humane use of animals in medicine.^[1–2]^ In addition, animal models are intrinsically low throughput and often fail to predict the efficacy and safety of drugs for human disease.^[3–6]^ These factors contribute to an expensive and inefficient drug development pipeline^[7]^ in which only 10% of drugs that enter clinical trials are ultimately approved.^[8]^ The success rate is even lower (8%) for drugs targeting central nervous system diseases.^[9]^ These diseases, including brain cancer, multiple sclerosis, Alzheimer’s Disease, and Parkinson’s comprised ∼ 14.7% of the global disease burden in 2020.^[10]^ These factors have motivated the development of *in vitro* models of human tissues collectively known as ‘tissue chips’ (also ‘microphysiological systems’ and ‘organs-on-a-chip’).^[11–12]^ With the advancement of some of these models, several tissue chip systems are already viable alternatives to animals for pre-clinical research.^[13]^ Combining these platforms with human induced pluripotent stem cell (hiPSC) technologies,^[14–15]^ tissue chip models are even being studied for their potential to simulate, and eventually contribute to, early stage human clinical trials.^[12]^ Importantly, tissues chips which require only small a volume of limited patient-derived materials and/or expensive therapeutics will be needed for these simulated clinical trials.

To realize their full impact on preclinical medicine, tissue chip platforms should become the preferred modality over conventional cell culture throughout the biomedical research community. The hesitancy to adopt tissue chip models in non-engineering laboratories can be traced largely to practical concerns including: 1) device and protocol complexity, 2) a lack of commercial accessibility, 3) low-throughput formats, and 4) missing reproducibility studies in expert laboratories. A related, but underappreciated, concern is the fact that pre-existing chip designs are often poorly suited to test a specific hypothesis or are incompatible with trusted assays. Often, acquiring the engineering resources and microfabrication skills for a custom tissue chip design is too high an entry barrier for many biomedical research laboratories.

We have previously introduced the µSiM as a Transwell™-style culture **micro**device featuring ultrathin (∼100 nm), highly permeable, and optically clear **si**licon nitride **m**embranes.^[16–18]^ We have used the µSiM to create *in vitro* models of the blood-brain barrier,^[19–21]^ the interface of the osteocyte lacuna-canalicular network in bone,^[22–24]^ and as a tool to study leukocyte transmigration across vascular endothelium.^[25–26]^ We addressed a need for high volume manufacturing and device reproducibility in a genetic screening of *Staphylococcus aureus* by enlisting a contract manufacturer,^[22, 24]^ but the approach produced large quantities of a configuration suitable to only one project. Many other ideas for µSiM-based tissue models require unique device configurations. The mass production of completed devices for each of these ideas is cost prohibitive, particularly during the discovery and validation phases of a project where design changes are expected and often necessary.

To maximize manufacturability and reproducibility while still enabling a design-flexible tissue chip platform, we now introduce the **m**odular **µSiM** (m-µSiM). The m-µSiM is modular in two senses: 1) <u>Modular assembly</u> allows the rapid construction from mass-produced components without requiring microfabrication tools or experience; 2) <u>Modular</u> <u>functionality</u> enables any component (membrane, upper well, bottom channel) to be quickly re-designed and replaced for a custom configuration that meets specific experimental needs. Modular functionality will also be apparent with forthcoming plug-and-play accessories that enable custom flow, measurements, and multiplexing without changing the core design. In this paper, we demonstrated the distribution and reproducibility of the m-µSiM through a collaboration between a bioengineering (University of Rochester, NY, “UR”) and a non-engineering brain barriers (University of Bern, Switzerland, “UniBe”) laboratories. Both laboratories demonstrated successful device assembly from components, the culture of an hiPSC-derived blood-brain barrier (BBB) model with low baseline permeability, and the expression of key junctional and adhesion molecules.

To support functional assessment of barrier function using small molecule permeability measurements, we developed and validated both an *in situ*-based method that takes advantage of the compatibility of the platform with high resolution microscopy, and a sampling-based permeability assay familiar to users of the conventional Transwell™ platform. We used the sampling assay in an interlaboratory reproducibility study that showed a statistically similar maturation of the BBB model over days in culture and statistical agreement with Transwell™ data in both laboratories. Finally, we demonstrated modular functionality through the introduction of a simple insert for side-by-side co-culture on two-membrane chips, and by exchanging nanoporous membranes for dual-scale, nanoporous/microporous membranes that enable leukocyte transmigration to the ‘tissue side’ of the platform. Modular functionality is most extensively demonstrated in a companion publication on a plug-and-play flow accessory for the µSiM.^[27]^ Importantly, these µSiM reconfigurations take minutes to assemble and require no knowledge of microfabrication techniques. Thus, by using mass produced components that easily ‘snap’ together, we have created a flexible design tissue chip platform accessible to any biomedical research laboratory.

## 2. Results

### 2.1. Design for Manufacturing with Rapid Local Assembly

The m-μSiM (**Figure 1**) comprises three mass-manufactured parts: 1) Component 1 is an acrylic block component featuring two fluidic ports for bottom channel access and a 100 µL well with a bottom ledge of exposed pressure sensitive adhesive (PSA) to enable sealing to the membrane chip; 2) Component 2 is an open bottom fluidic channel of ∼10 µL in total volume. It is made from stacked PSA/polyethylene terephthalate (PET) layers with a 50 µm cycloolefin polymer (COP) bottom layer that provides glass-like optical clarity. The top of Component 2 is PSA that enables bonding to Component 1; 3) The third part is the membrane ‘chip’ which can be selected from nanoporous, microporous, or dual-scale (a mix of micro and nanopores) options based on application needs (Figure 1A, C).^[18–20]^ Membrane chips have a trench side and flat side containing the membrane.^[25]^ The membrane is free-standing over a window of 700 µm x 2 mm. In this study, chips are oriented “trench-down” to culture a monolayer that is continuous across the window and non-window regions, however, higher resolution imaging can be achieved by flipping the chip into a “trench-up” orientation to bring the cells closer to the objective lens (Table S1, Supporting Information).^[20]^

**Figure 1.**
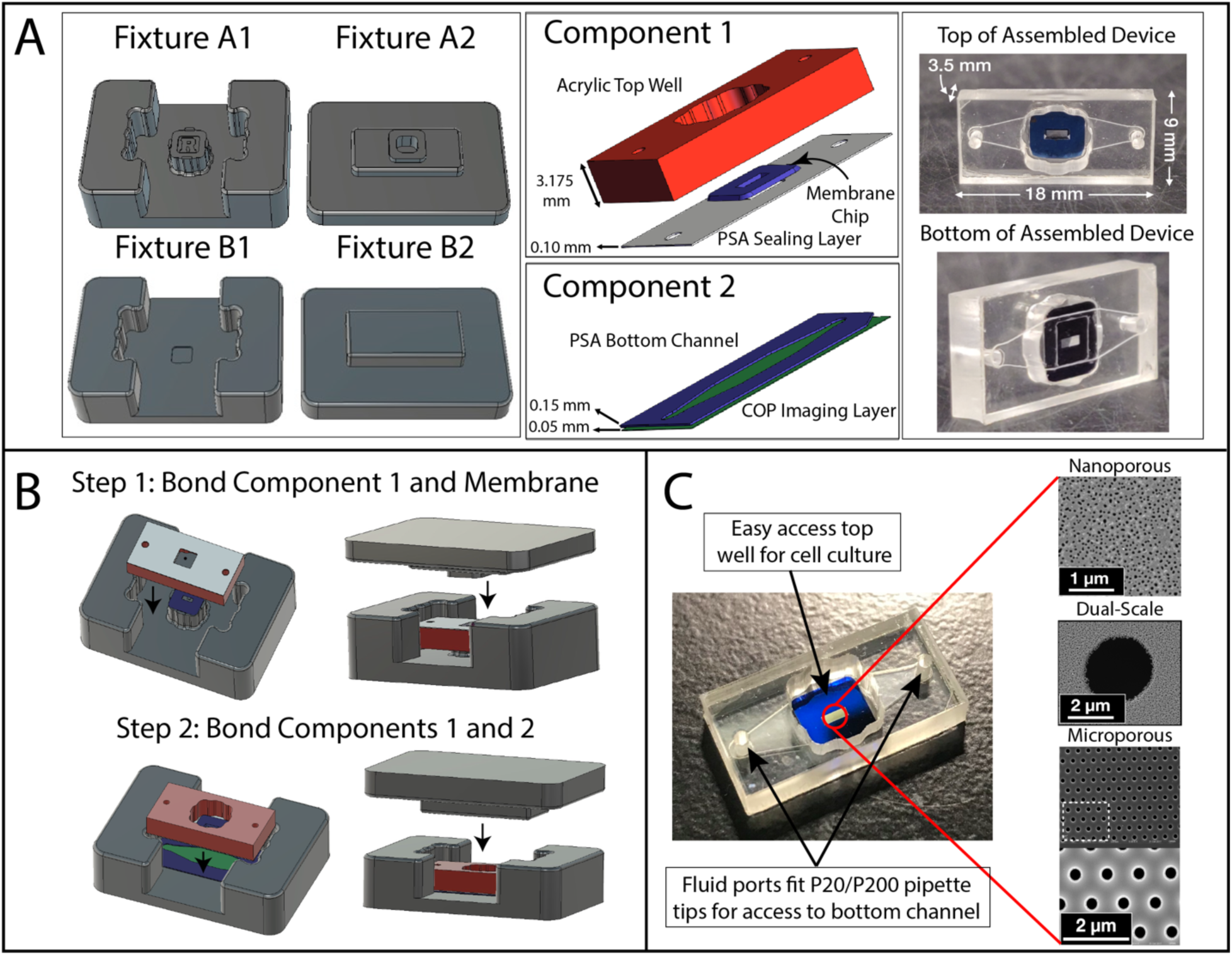
m-µSiM assembly. (A) Fixtures are used to guide components and membrane chip together (left). Component 1 is composed of an acrylic top layer with a Transwell^TM^-style open well and a PSA sealing layer. Component 2 is composed of a thin bottom channel and COP imaging layer (middle). Assembled devices are 18 x 9 x 3.5 mm. Displayed is a “trench up”-style device (right). (B) Assembly is a two-step process. *Step 1: Bond Component 1 and membrane.* The membrane chip is placed on Fixture A1 pedestal. Place Component 1 face-down over membrane. Use Fixture A2 to press firmly and activate PSA. This irreversibly bonds membrane to Component 1. *Step 2: Bond Components 1 and 2.* Place Component 2 in Fixture B1, channel-side up. Place Component 1 with the membrane chip onto Component 2. Use Fixture B2 to press firmly to activate PSA, irreversibly bonding Component 1 and Component 2. (C) The modular assembly allows for easy reconfiguration for the application at hand. The example here illustrates the choice of different membrane architectures. The device displayed is a “trench-down”-style device. Component 1’s open well format allows easy cell culture, and access ports provide access to the bottom channel. They are designed to seal-to-fit standard P20 and P200 pipette tips.

Assembly of the modular µSiM is a two-step process that utilizes PSA to irreversibly bond all components together (Video S1, Supporting Information). The PSA layers are protected by removable liners, which are added during component manufacture and removed just prior to m-µSiM assembly. A pair of two-piece fixtures are used to press components together in alignment (Figure 1B). In step one, the membrane chip is placed on Fixture A1 in the opposite orientation to the desired final orientation (“trench-up” or “trench-down”). Component 1 is placed over it in an inverted fashion, and Fixture A2 is used to apply pressure to the backside of Component 1. This affixes the membrane chip to Component 1, as the non-porous boundary of the chip bonds to the PSA sealing layer at the bottom of the well. In step two, Component 2 is first placed into Fixture B1 with the channel-side facing up. Component 1 with the membrane chip is placed on top of Component 2, and Fixture B2 is used to apply pressure and bond the two components together. Assembly time is under 5 minutes, compared to the many hours required for traditional, UV ozone-based assembly. Further, a dye leak test confirmed proper sealing of devices, and a LIVE/DEAD stain on the commercially-available human brain microvascular endothelial cell line, hCMEC/D3, established basic biocompatibility (Figure S1, Supporting Information).

### 2.2. *In Situ* Measurements of Small Molecule Permeability

Control of small molecule permeability is a primary function of vascular barriers. Unlike peripheral microvascular beds, which allow for diffusion of small molecules, the BBB is a highly restrictive barrier that is impermeable to most small molecules, with healthy barriers allowing access only to molecules with carriers or transporters.^[28]^ Thus, we developed two assays to measure barrier permeability to small molecules in the m-µSiM, both of which were designed for measurements in the small volume (∼10 µL) of the ‘receiver’ channel of Component 2 (tissue-side) when dye is added to the donor well of Component 1 (blood-side). The first assay involves use of confocal microscopy to directly image the evolution of fluorescence in the trench of the membrane chip directly beneath the endothelial barrier after addition of a fluorescence tracer to Component 1’s well (**Figure 2A**). This assay takes advantage of the compatibility of the m-µSiM with inverted live cell microscopy to make non-invasive measurements of small molecule permeability *in situ*.

**Figure 2.**
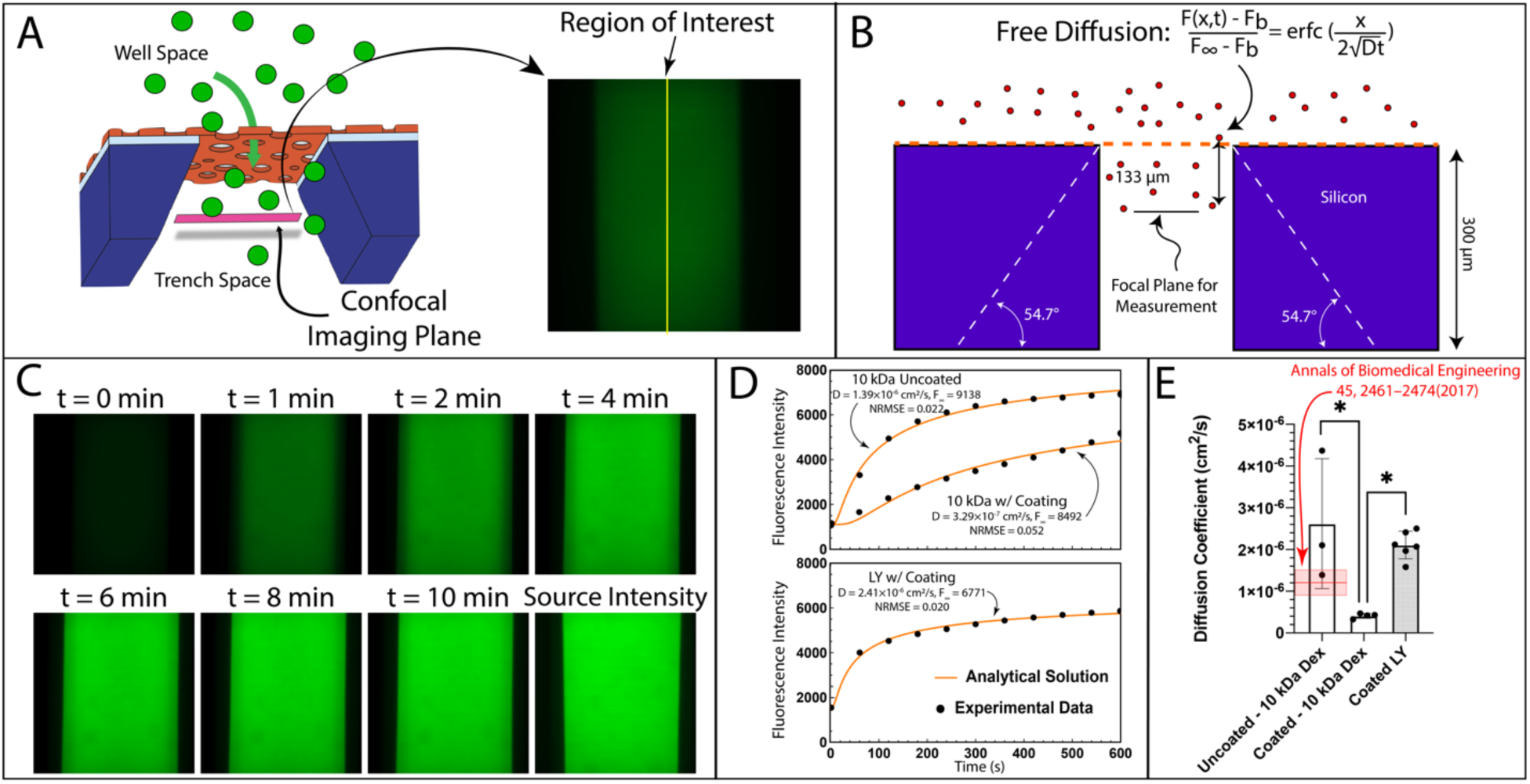
*in situ* permeability assay optimization on cell-free devices. (A) The imaging plane of a confocal microscope is focused 133 µm below the membrane, within the chip’s trench. Dye diffuses from the well into the trench (left). An example of corresponding image (right) shows a linear region of interest in the center of the membrane (yellow line) where 1-D diffusion accurately describes the evolution of fluorescence (see Figure S2, Supplemental Information). (B) The diffusion coefficient, ***D***, and fluorescence as time reaches infinity, ***F_∞_***, are solved using 1-D Fick’s Law describing free diffusion. (C) Representative images illustrating 10 kDa Dextran-AF488 diffusion into the trench of an uncoated membrane chip over the course of ten minutes. Following the ten minute diffusion, dye from the top well was flushed across the bottom channel to obtain a “Source Intensity” photo. (D) Example plots of diffusion across uncoated and collagen/fibronectin-coated chips using 10 kDa Dextran-AF488 (top) and lucifer yellow (LY, bottom). The analytical solutions fit well to the experimental data (normalized root mean square error, NRMSE, < 0.1). (E) The resulting diffusion coefficients for 10 kDa Dextran-AF488 across uncoated and collagen/fibronectin-coated membranes and LY across collagen/fibronectin-coated membranes. Coating the membrane significantly decreased the apparent diffusion coefficient and larger diffusion coefficients are measured for the smaller molecule, LY. The rapid diffusion through an uncoated membrane was challenging to measure but confirmed a negligible hindrance of the dye by the membrane compared to coated membranes. It overlapped with fluorescence correlation spectroscopy measurements (red bar). N = 3-6 per group. Ordinary one-way ANOVA, p < 0.05.

Unlike conventional clearance measurements for barrier cells, our *in situ* measurements of permeability are not taken at steady state. Thus, we sought to interpret time-dependent rise of fluorescence intensity using a 1-D analytical solutions of transport into a semi-infinite space. One concern was the possibility that the sloping walls of the silicon ‘trench’ on the backside of membrane chips,^[25]^ would preclude the use of a 1-D analytical equation (Figure 2B). To address this, we accurately modeled the trench as a 2-D geometry in COMSOL Multiphysics and compared 2-D simulations of diffusion into the trench to the 1-D analytical model. We found no differences in the temporal evolution of concentrations at the centerline, 100 µm below the membrane for a range of relevant diffusion coefficients representing < 1 kDa; 10 kDa; and 40 kDa tracers (Figure S2, Supporting Information). We conclude that for measurements taken at the center of the membrane, the influence of the sloping trench walls can be ignored.

We then measured the free diffusion of both Alexa-488 tagged 10 kDa Dextran (10 kDa Dextran-AF488) and lucifer yellow (LY, 457 Da) across uncoated and collagen/fibronectin-coated membranes to validate our experimental set up. Coating was necessary to slow diffusion across our highly porous membranes. In these experiments we first focused on the membrane and then precisely lowered the objective to a position 100 µm beneath the membrane. It is important to note, that while the objective physically moves 100 µm, the index mismatch between air and media at the sample surface results in a focal plane shift of studies with non-porous membranes established that this was the minimum position where background measurements became independent of the location of z-focus (Figure S3B, Supporting Information). Preliminary studies were also done with each molecule to establish that the optimized exposure times resulted in no detectable photobleaching or other instability during image acquisition (Figure S3C-D, Supporting Information).

These cell-free experiments were initiated by replacing the full m-µSiM well volume (100 µL) with a solution containing the optimized concentration of tracer dye. An imaging sequence was initiated within 3 seconds of adding the tracer, with images taken every minute for 10 minutes (Figure 2C). Centerline values were extracted from images of the membrane (Figure 2A, yellow line). As exemplified by the curves in Figure 2D, Equation (1) provided quality fits to data, indicating that free diffusion was a good description of the physics governing the evolution of fluorescence in these experiments. Importantly, we were able to detect significantly slower diffusion of 10 kDa Dextran-AF488 through collagen/fibronectin-coated membranes compared to the small dye LY (457 Da) (Figure 2E). In the case of an uncoated membrane, while our results were high variable, the measured diffusion coefficient overlaps with published values obtained by fluorescence correlation spectroscopy (FCS) measurements.^[29]^ It is possible that on an uncoated membrane, small pressure differences across the membrane lead to convective transport. However, for coated membranes, for which transport was slower and convective contributions are likely less of a concern, we obtained much more consistent results. Importantly, the clear difference between coated and uncoated membranes with 10 kDa Dextran-AF488 in these experiments affirms a key value proposition for the use of ultrathin membranes in tissue chip models: because of their thinness, the membranes themselves provide no practical hindrance to the diffusion of molecules significantly smaller than the pores of the membranes. We have shown this before in several detailed studies of membrane transport by diffusion.^[16, 30–31]^ Thus, for tissue chip models created with ultrathin membranes, only the biological barriers derived from cells and extracellular matrices will determine the rate of small molecular exchange between compartments.

Having validated our *in situ* approach with computational, analytical and experimental studies of free diffusion, we turned to cell permeability measurements (**Figure 3**). The 1-D analytical form to interpret these studies is the solution to molecular transport into a semi-infinite space from a boundary experiencing constant flux (see Equation (6), Methods) (Figure 3B).^[32–33]^

**Figure 3.**
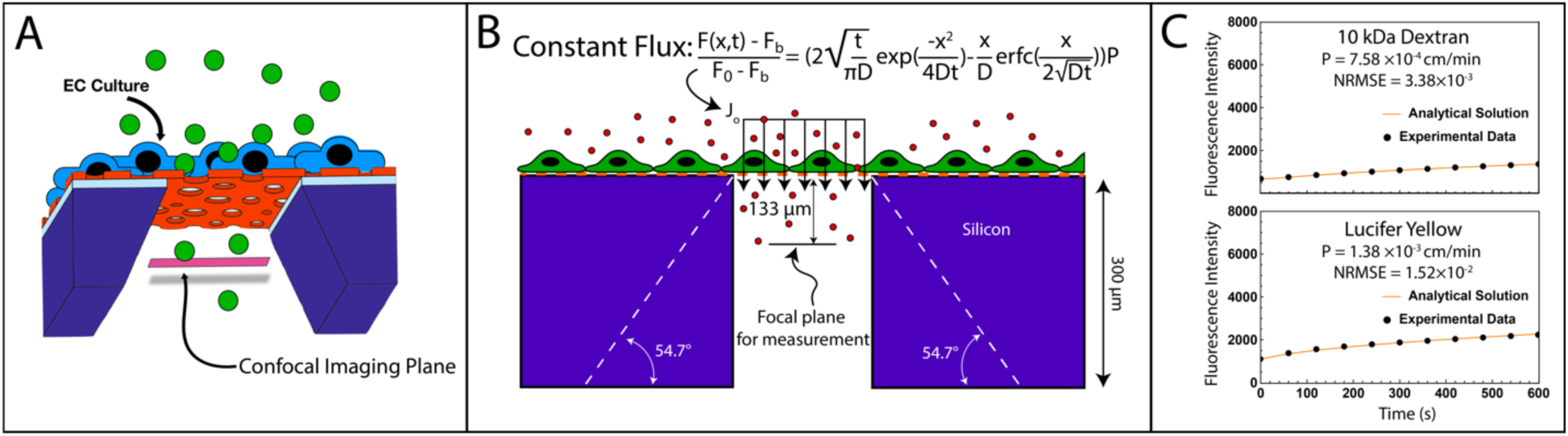
*in situ* permeability assay optimization using hCMEC/D3. (A) A confocal microscope is focused 133 µm below the membrane, within the chip’s trench. Dye diffuses from the well, across an endothelial cell layer, and into the trench. (B) Endothelial permeability is calculated assuming constant flux across the monolayer (see Methods). (C) Example plots of the analytical solutions for permeability of 10 kDa Dextran-FITC (top) and 457 Da lucifer yellow (bottom) across an hCMEC/D3 monolayer. The analytical solutions fit well to the experimental data, with low normalized root mean square errors (NRMSE, < 0.1).

Example plots for *in situ* permeability experiments with hCMEC/D3 cells are shown in Figure 3C for both 10 kDa Dextran conjugated to FITC (10 kDa Dextran-FITC) and LY. While an AF488 conjugation is preferred for the *in situ* assay due to superior photostability and FITC’s known pH-sensitivity,^[34]^ we could only find literature sources to benchmark against for FITC-labeled dextran. Both tracers showed transport to the measurement plane was significantly slowed by the presence of the monolayer compared to experiments without cells, with the smaller LY showing a more rapid increase in fluorescence on the same 10-minute time scale compared to 10 kDa dextran. Equation (6) provides excellent fits to the data (normalized root mean square error < 0.1) with permeability values that lie within the range found in the literature (see summary panel for both methods **Figure 4E**).

**Figure 4.**
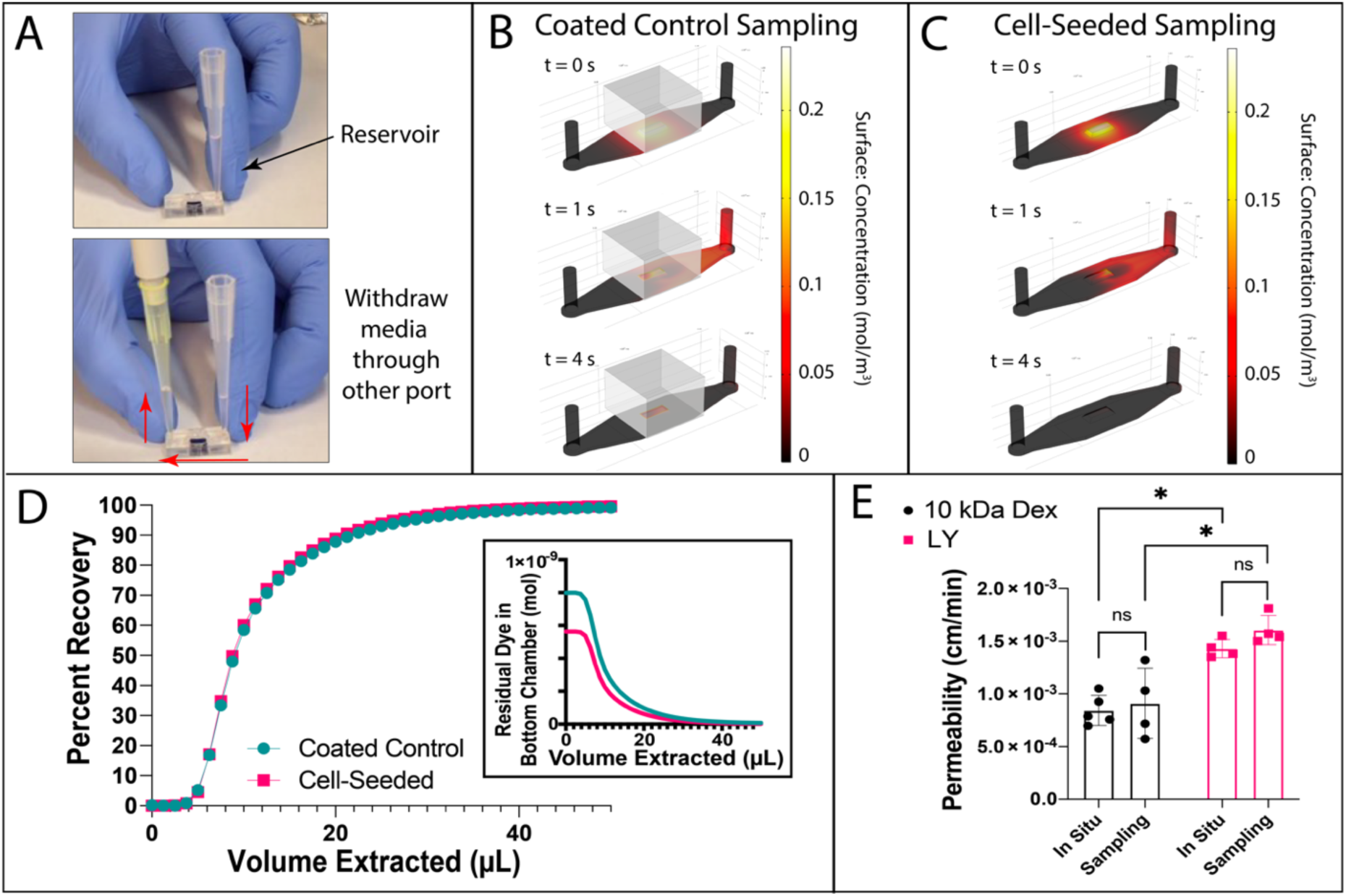
Sampling permeability assay optimization and comparison to *in situ* method. (A) Illustration of the sampling method for collecting dye from the channel. A reservoir pipette tip is added to one port that accesses the bottom channel, and another pipette tip is used to pull media out via reverse pipetting. Media withdrawn is added to a well plate for fluorescence measurements. COMSOL Multiphysics was used to model the diffusion (B) or flux (C) of lucifer yellow across coated control (B) or cell-seeded devices (C). Time zero illustrates the dye in the bottom channel after one hour of diffusion or flux, prior to flushing the solution out of the channel. t = 1 second illustrates the flushing process across the channel and out the right port, and t = 4 seconds shows remaining dye after the flushing is complete. (D) COMSOL-generated data was used to determine the volume needed to clear analyte from the channel. The plot shows percent recovery of transported dye, defined as the ratio of total analytes extracted to the total transported, and residual dye left in the chamber in moles (inset). (E) hCMEC/D3 permeability to 10 kDa Dextran-FITC (10 kDa Dex) and lucifer yellow (LY). There were no significant differences in measured permeability between the *in situ* and sampling methods for either dye. Both methods measured significantly higher permeability of hCMEC/D3 to lucifer yellow (LY) compared to 10 kDa Dextran-FITC (10 kDa-Dex). N = 4-5 per group. Two-way ANOVA with Tukey *post hoc* test, p < 0.05.

A subtle but notable difference between the *in situ* method and traditional methods for permeability measurements is the lack of a coated membrane control for calculating endothelial permeability. In traditional assays, a coated membrane control accounts for the diffusive resistances of the membrane, the membrane coating, and the device geometry (or ‘system’). However, the equivalent permeability of the coated membranes is >> 1 cm/min (using P_e_ ∼ D/L as estimate of the permeability of the coated membrane where L is 0.1 µm for the membrane thickness and D is the diffusion coefficient measured across a coated membrane; Figure 2E). Because this value is four orders of magnitude higher than cell permeabilities, subtracting the hindrance caused by a coated membrane would have no impact on the results. Note that the lack of a system permeability subtraction step is a direct benefit of the ultrathin membrane technology used in the µSiM. While not directly tested here, the method should improve the ability to detect changes in monolayer permeability in response to inflammatory signals and similar influences.

### 2.3. Sampling (Endpoint) Assays of Small Molecule Permeability

The second assay we developed is an analog of the conventional method for small molecule permeability in Transwells™ that samples the dye that has cleared from the top to the bottom compartment over time.^[35]^ We developed this as a more accessible approach to permeability measurements in the m-µSiM, as it does not require the careful use of confocal microscopy and the theory and calculations should be familiar to laboratories who already do Transwell™-style permeability measurements. The key innovation for this protocol was devising a method to fully sample the contents of the bottom channel despite the fact that it only contains 10 µL of total fluid and is not actively mixed. We achieved this by adding a 50 µL pipette “reservoir” with blank media to one port and ‘reverse pipetting’ a 50 µL volume with a pipette attached to the other port (Figure 4A; Video S2, Supporting Information). Because the full volume is harvested in this assay, and because the impact of complete basal media exchange on the monolayer above is unknown, we considered our approach to be an ‘end-point’ assay. Thus, unlike traditional Transwell™ studies, which sample from the basal compartment repeatedly over time, we sample exactly once at the end of an hour-long incubation with the dye.

Because of the complex geometry of the bottom channel including the chip trench, we used COMSOL Multiphysics to check our expectation that flushing this space with clean media should harvest all of the tracer molecules that pass through the monolayer and membrane. We first modeled the m-µSiM geometry (chip + Component 2) with both free diffusion and constant flux boundary conditions at the membrane over an hour of transport into the receiving channel. We then simulated flushing and collecting media from the backside channel (Figure 4B-C). The simulated concentration profile after 1 hour of permeation and lateral diffusion is shown at t = 0 seconds. At t = 1 second and t = 4 seconds, the analyte concentration profile change during the 50 µL sampling process is illustrated. Analyzing the extracted volume allowed us to calculate the expected amount of residual analyte that remains in the backchannel as a function of the volume flushed through the device (Figure 4D, inset). Normalizing this data to a ‘percent recovery’ (see Methods) shows that regardless of the method of tracer transport, a 20 µL flush volume is expected to remove ∼88-89% of the analyte from the backside channel, a 40 µL flush volume will remove ∼98-99% and a 50 µL flush volume will remove over 99% of the tracer molecule (Figure 4D). Percent recovery using a 50 µl flush volume was validated experimentally, finding ∼97% dye recovery (Figure S3E, Supporting Information), which we consider acceptable. Once collected, monolayer permeability can be calculated with methods and equations used routinely for Transwells™.^[35]^ Note that as part of this classic methodology, the “system permeability” must also be measured using a cell-free, coated control and subtracted to determine the cellular contribution (see Methods).

With both the *in situ* and sampling-based methods of measuring endothelial permeability established, we compared results between the two assays for the two tracer molecules across hCMEC/D3 monolayers (Figure 4E). Despite the *in situ* assay not needing a coated-control, the two methods gave agreement in permeability measurements for both 10 kDa dextran and LY, with each detecting significantly higher permeability for the smaller LY. Importantly, the permeability values for both dyes fall within the expected, albeit large, ranges based on reported literature for both 10 kDa Dextran-FITC (0.84 ± 0.14 x 10^-3^ cm/min *in situ*, 0.91 ± 0.33 x 10^-3^ cm/min sampling, 0.28 – 2.04 x 10^-3^ cm/min literature^[36–38]^) and LY (1.43 ± 0.09 x 10^-3^ cm/min *in situ*, 1.61 ± 0.14 x 10^-3^ cm/min sampling, 0.60 – 1.55 x 10^-3^ cm/min literature^[39–41]^).

### 2.4. Establishment of the EECM-BMEC-like Cell Culture on the m-µSiM

Next, we sought to establish the reliability of the m-µSiM for use in a non-engineering brain barriers laboratory (**Figure 5**). In these experiments, the UniBe lab made use of their recently developed stem-cell based model of the BBB by differentiating hiPSCs into extended endothelial culture method (EECM) brain microvascular endothelial cell (BMEC)-like cells^[42–43]^ and cultured the cells in the m-µSiM. To confirm proper barrier maturation, EECM-BMEC-like cells were stained for adhesion and junctional complex molecules. Images were acquired at 10X magnification and zoomed in digitally to quarter-size. (Figure 5A). EECM-BMEC-like cells cultured in m-µSiMs expressed key BMEC junctional molecules (Figure 5B). Further, EECM-BMEC-like cells cultured in the m-µSiM showed a clear upregulation of leukocyte adhesion molecules (ICAM-1; VCAM-1) after stimulation with proinflammatory cytokines (TNFα+INFγ) and constitutive expression of ICAM-2, replicating the published findings for EECM-BMEC-like cells cultured on Chamber Slides^TM^ and Transwell^TM^ filter inserts (Figure 5C).^[43]^ The full set of stains shown in Figure 5 was reproduced in the bioengineering laboratory at UR using EECM-BMEC-like cells independently differentiated from the same clonal line of hiPSCs (Figure S4, Supporting Information). These results establish the successful exchange and reproducibility of numerous m-µSiM-specific protocols even when employing sophisticated brain barrier models. The reproduced protocols include device assembly, membrane coating, cell seeding and maintenance to barrier maturation, fixation, and staining.

**Figure 5.**
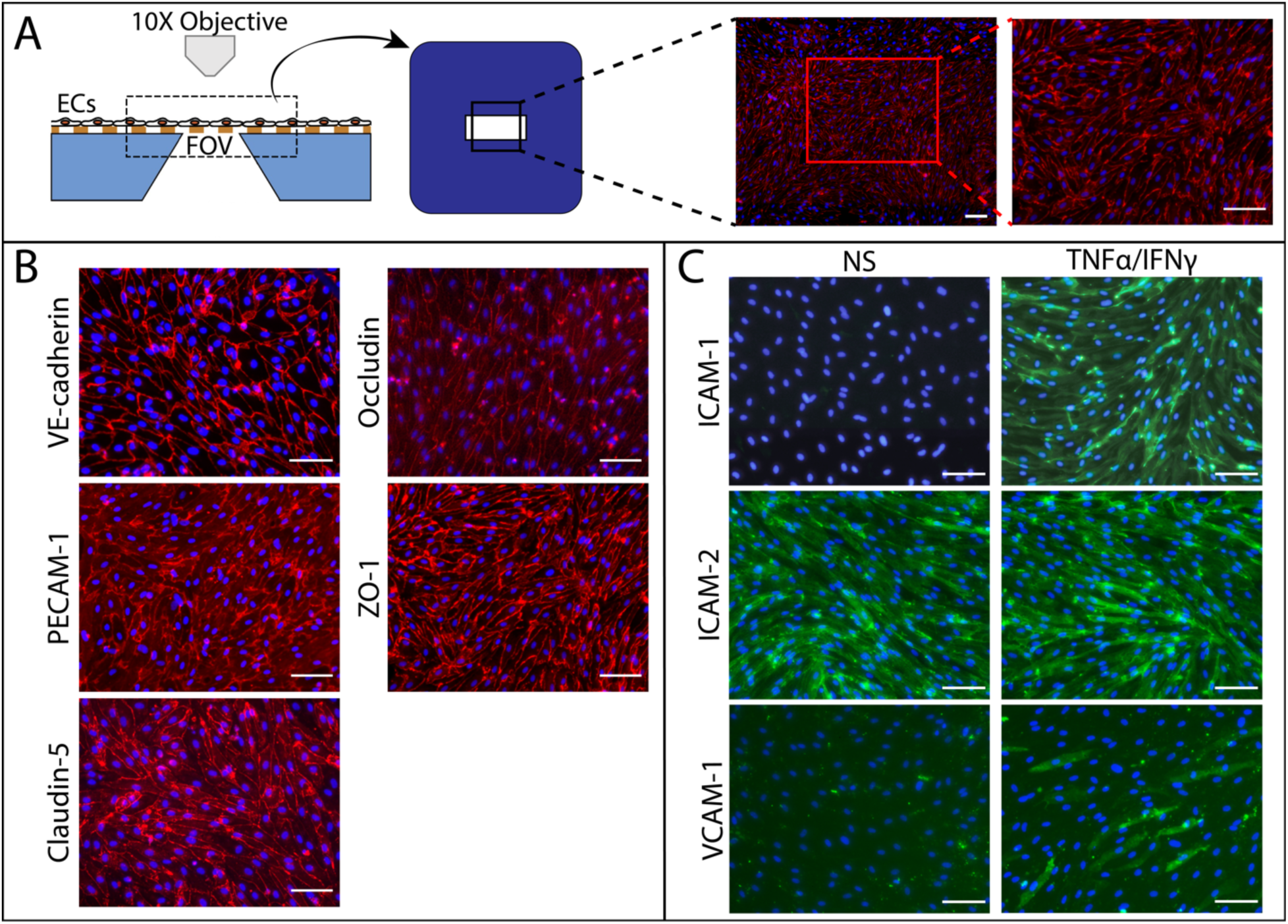
Establishment of EECM-BMEC-like cell culture in the m-µSiM by a brain barriers laboratory. (A) Images of endothelial cells (ECs) at UniBe were acquired on a Nikon E600 Fluorescence microscope using a 10X objective. The field of view (FOV) was centered on the membrane. Images were digitally cropped post-acquisition to better visualize molecular stains. (B) EECM-BMEC-like cells expressed key junctional molecules when cultured on m-µSiM devices. (C) EECM-BMEC-like cells expressed key cell adhesion molecules upon exposure to proinflammatory stimuli when cultured on m-µSiM devices. Non-stimulated (NS) and stimulated (0.1 ng/ml TNFα + 2 IU/ml IFNγ) for 16-20 hours. Cells have comparable expression patterns to published data from Chamber Slides^TM^ and Transwells^TM^. Scalebar = 100 µm.

### 2.5. Interlaboratory Reproducibility of EECM-BMEC-like Cell Permeability

To quantitatively assess the interlaboratory reproducibility of establishing a barrier model on the m-µSiM, we used the sampling-based permeability assay with LY to evaluate EECM-BMEC-like cell barrier maturation over time at both UR and UniBe (**Figure 6**). We also compared the results on the m-µSiM to conventional Transwell™ cultures. Both labs saw high variability in the permeability of monolayers cultured for only 2 days in the m-µSiM, underscoring that 2 days of culture are insufficient for barrier maturation. After 4 days, UniBe was able to achieve reproducible permeability values that matched Transwell™ data to sodium fluorescein from the original EECM-BMEC-like cell publication, differentiated from the same hiPSC clone, IMR90-4 (m-µSiM: 0.60 ± 0.08 x 10^-3^ cm/min, published Transwell™: ∼0.65 x 10^-3^ cm/min).^[43]^ At UR the 2, 4, and 6-day cultures in the m-µSiM steadily matured and improved reproducibility over time. Permeability values from the UR study were statistically indistinguishable from the corresponding timepoints at UniBe. Moreover, after 6 days of culture in the m-µSiM, the UR studies agreed with their 6 day Transwell™ data (m-µSiM: 0.52 ± 0.01 x 10^-3^ cm/min, Transwell™: 0.40 ± 0.06 x 10^-3^ cm/min) suggesting no detectable difference in barrier function between the two platforms. Thus, these studies quantitatively demonstrate that m-µSiM assembly, culture, and sampling-based permeability can be reproduced between two labs, even with a sophisticated hiPSC-based BMEC model. Furthermore, the data also demonstrate that we can achieve comparable barriers to LY on our platform to those formed on conventional Transwells™, matching the current ‘gold-standard’ assay.

**Figure 6.**
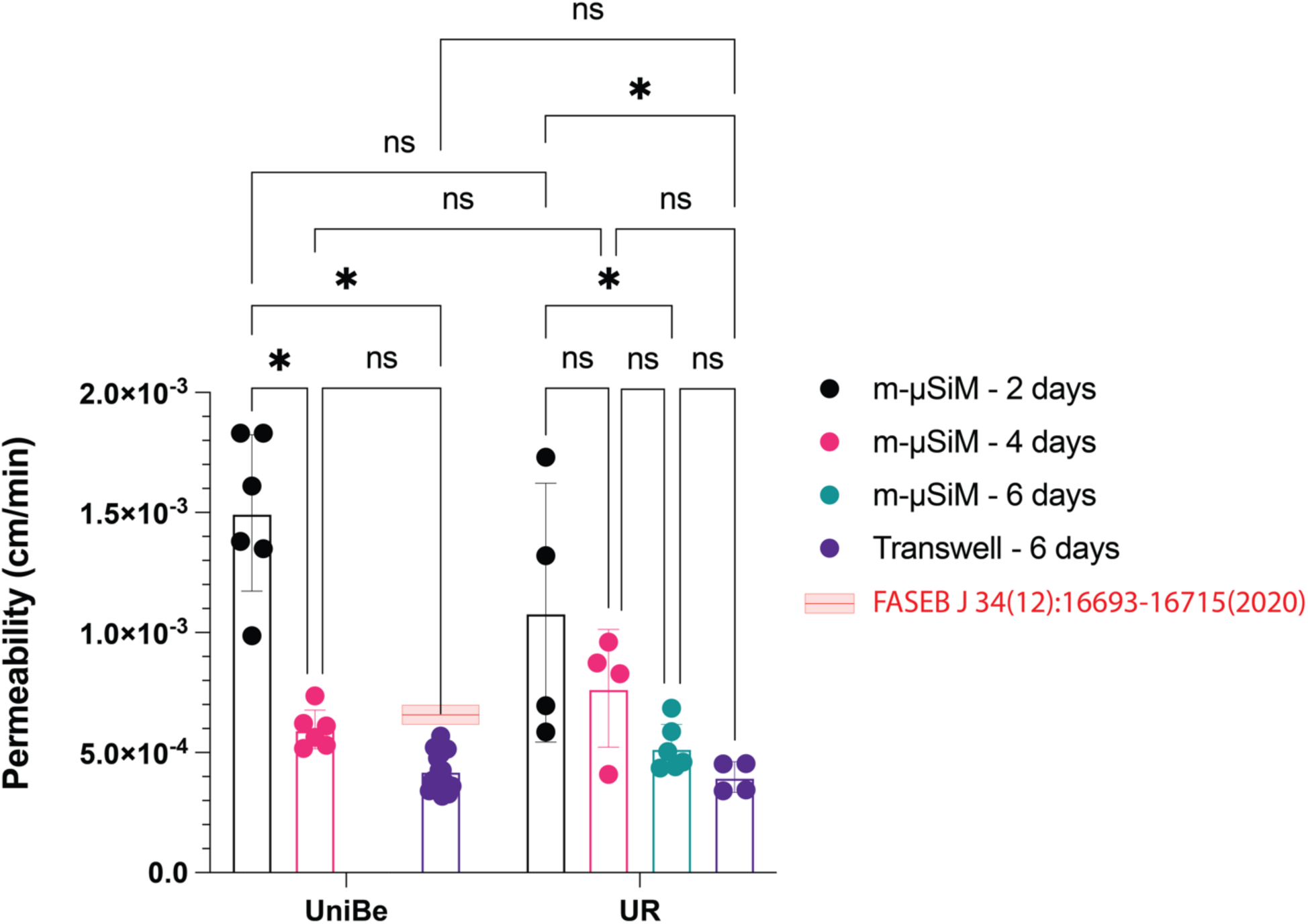
Interlaboratory reproducibility of EECM-BMEC-like cell permeability on m-µSiM between non-engineering brain barriers and bioengineering laboratories. EECM-BMEC-like cells were cultured in the m-µSiM for 2, 4, or 6 days or in Transwell^TM^ filters for 6 days, and permeability was measured using the sampling assay (m-µSiM) or clearance assay (Transwell^TM^) at both the University of Bern (UniBe) and University of Rochester (UR). There were no significant differences in permeability between labs or culturing platforms upon cell maturation (4-6 days in m-µSiM). Data was comparable to UniBe’s previously published data of EECM-BMEC-like cells cultured in Transwells^TM^ and assayed for permeability to sodium fluorescein (red bar, FASEB J 34(12):16693-16715(2020)). N = 4-16 per group. Two-way ANOVA to compare labs and culturing conditions, excluding m-µSiM – 6 days with Tukey *post hoc* test; one-way ANOVA to compare within UR culturing conditions, p < 0.05.

### 2.6. Demonstration of Modular Functions

Our final studies demonstrate with a few brief examples the ability of the m-µSiM platform to be quickly reconfigured to address different experimental needs with few brief examples (**Figure 7**). An extensive example of modular functionality is given in a companion paper by Mansouri et al. describing the development and use of a flow module to introduce controlled fluid flow in the m-µSiM.^[27]^

**Figure 7.**
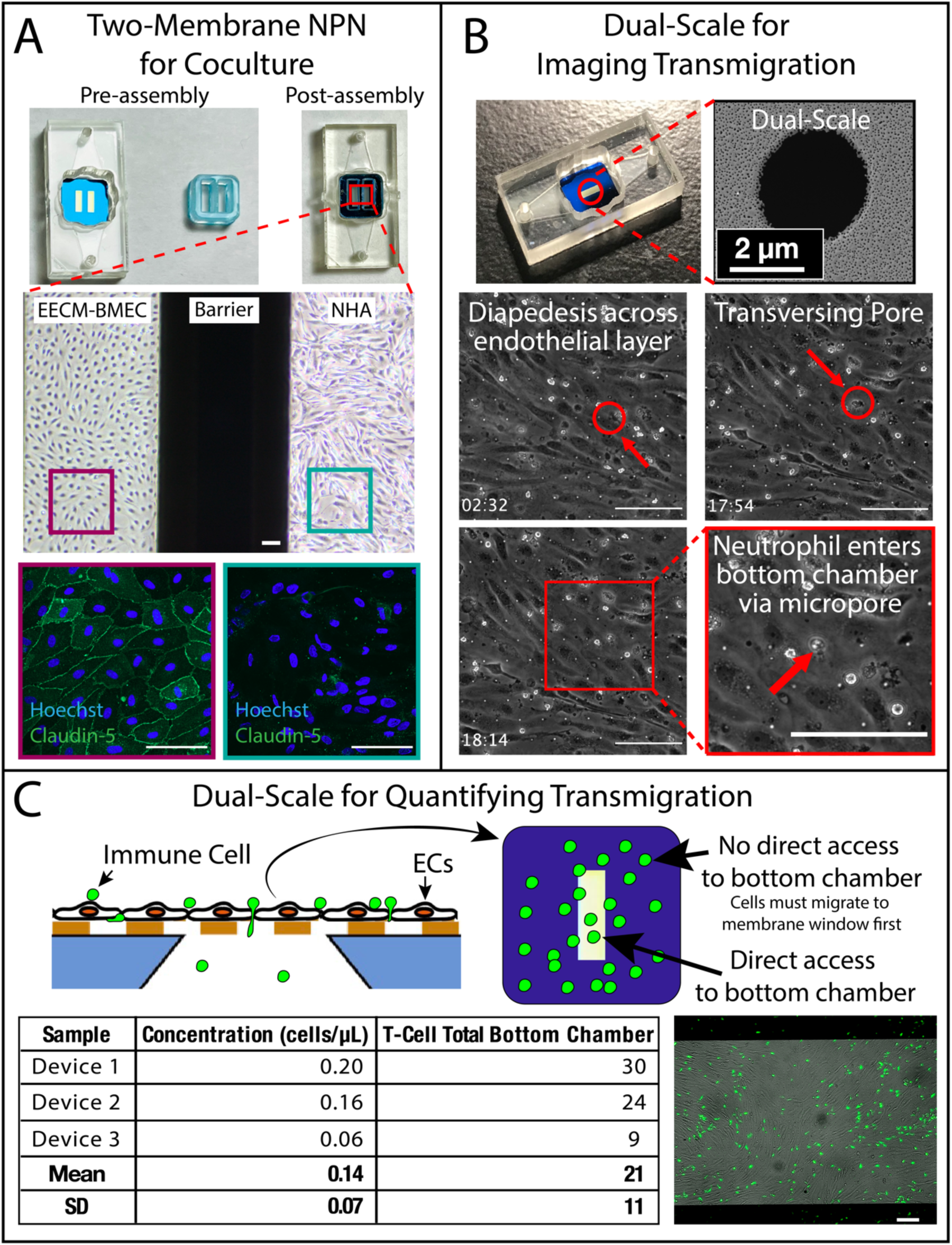
Demonstration of the modular function of m-µSiM. (A) Side-by-side co-culture was achieved by swapping one window nanoporous (NPN) membranes with two window NPN membranes and by addition of a cell culture insert. EECM-BMEC-like cells were cultured in one chamber and human astrocytes (NHA) in the other chamber, with no apparent cross-contamination of cells. Chambers were stained for EECM-BMEC-like cell marker, Claudin-5 (green) and nuclear marker Hoechst (blue). Phase images were acquired on a Nikon Eclipse Ts2 phase contrast microscope and fluorescence images on an Andor Spinning Disc Confocal Microscope. Scalebar = 100 µm. (B) Neutrophil migration across EECM-BMEC-like cells cultured on 0.625%, 3 µm dual-scale membranes. Neutrophils were seen migrating across the endothelium (02:32) and through a micropore (17:54), entering the bottom channel (18:14) (time in min:sec). Videos were acquired on a Nikon Ti2 Eclipse inverted microscope using a long working distance 40X objective in phase contrast. (C) T-cell migration across 1.25%, 3 µm dual-scale m-µSiM was quantified by flow cytometry of CellTracker™ Green CMFDA-labelled migrated T-cells. Transmigrated T-cells can only access the bottom channel at the membrane window region of the chip. Remaining adhered T-cells were visualized via epifluorescence imaging, paired with phase contrast imaging of the endothelial layer. Images were acquired on a Nikon Eclipse E600 Microscope using a 10X objective. Scalebars = 100 µm.

The first example involved two changes to the standard m-µSiM to establish a side-by-side co-culture that limits paracrine signaling to the basal chamber. While astrocytes and BMECs are often cultured on opposite sides of a membrane to model the neurovascular unit as a proximal co-culture, a side-by-side configuration could be used to study paracrine signaling in an indirect co-culture with the added benefit that both cell types can be clearly imaged because they are not in the same optical path. To achieve this, the conventional single membrane chip was replaced with a two-membrane chip, and a cell culture chamber insert was placed in the well to create side-by-side chambers such that one membrane is in each chamber (Figure 7A). We conducted dye leak tests to establish that the only path for small molecule exchange between the Component 1 chambers is through their common channel-side compartment (Figure S5, Supporting Information). We demonstrated that EECM-BMEC-like cells seeded in one of the resulting sub-chambers of the m-µSiM well and primary human astrocytes seeded into other chamber resulted in no apparent cross-contamination of cells (Figure 7A; Figure S6).

As a second example, m-µSiMs were built using ‘dual-scale’ nanomembranes, in which micropores are etched onto a nanoporous background.^[18]^ The nanoporous membranes (∼60 nm diameter pores) used thus far allow diapedesis across the endothelial layer but prevent immune cell entry into the bottom channel.^[26]^ Dual-scale membranes feature additional 3 µm pores patterned at a low enough density (0.625-1.25% additional porosity) so as to not compromise imaging. While the dual-scale membranes do not obscure imaging, we did find that the meniscus from the open well of the m-µSiM did compromise live-cell imaging by phase contrast microscopy. This was solved by addition of the flow module insert described in our companion paper to create a closed chamber, and injecting neutrophils into one of the module’s ports.^[27]^ Figure 7B demonstrates high resolution, phase contrast imaging of neutrophil transmigration across an EECM-BMEC-like cell monolayer after a potent neutrophil chemokine is added to the bottom chamber (Video S3, Supporting Information). The image quality matches those seen with preceding flow-cell devices that were assembled via a labor-intensive ozone plasma approach by our group.^[18]^

A final example also uses dual-scale membranes for an assay of T-cell migration into the basal chamber of the m-µSiM, quantified by flow cytometry (Figure 7C; Figure S7, Supporting Information). Because this experiment did not require high resolution imaging, injection via the flow module was not necessary. Instead, T-cells were fluorescently-labeled with CMFDA and directly added to the open well of the m-µSiM above an EECM-BMEC-like cell monolayer. T-cells that migrated spontaneously into the bottom chamber were collected after a two-hour incubation using the same backchannel flushing technique developed for sampling permeability studies (Figure 4A) and quantified using flow cytometry. One important note about this experiment is that immune cell access to the bottom chamber is limited to the active membrane region of the chip, which accounts for ∼5% of the total surface area of the monolayer. Despite this limitation, the m-µSiM enables the quick observation and imaging of T-cell interactions with the monolayer, whereas Transwell™ protocols require tedious fixation, cutting out filters, and staining to visualize the monolayer and count adhered immune cells. As shown in Figure 7C, the green fluorescent T-cells can be seen in the same image as the endothelial monolayer by simply acquiring a phase contrast and epifluorescence image of the culture after quick fixation and washing.

## 3. Discussion

The past decade has seen a dramatic rise in tissue chip development and application.^[11]^ For vascular barrier models, in particular, there are now many examples ranging from simple to highly complex.^[38, 44–47]^ Sophisticated microdevices are often difficult to use outside of the engineering laboratories that create them, making it challenging to establish confidence in a model through interlaboratory reproducibility. While mass-produced commercial platforms overcome concerns with device reproducibility with relatively simple designs, they also limit users to a particular geometry, which is likely to be sub-optimal for the assays, hypotheses, and/or tissue model of interest. For example, the popular tissue chip platform produced by Emulate is widely used by the research community but is a fixed design.^[48]^

To address the need for design and assay flexibility, the concept of ‘plug-and-play’ modularity in tissue chip platforms has recently emerged.^[49–50]^ Our modular version of the established µSiM platform provides non-engineering (and engineering) labs with an accessible and adaptable option for modeling tissue barriers. We demonstrated facile assembly and reproducibility through a collaboration between physically distant laboratories (UniBe, Switzerland and UR, USA). Lack of data reproducibility is not unique to the tissue chip field,^[51]^ but may be exacerbated by the complexity of tissue chip devices.^[52]^ Despite these challenges, both laboratories established EECM-BMEC-like cell cultures from the commercial IMR90-4 hiPSC clonal line^[42–43]^ and showed that they exhibit key characteristics of brain microvascular endothelial cells in the m-µSiM,. Underlying this agreement between distant laboratories are numerous jointly-established protocols for membrane coating, cell seeding, culture maintenance, fixation, immunofluorescence staining, imaging, and more. To equip future users, consensus protocols have been developed and are now ready for further dissemination through this publication and through freely accessible web pages (https://nanomembranes.org/resources/modular-μsim/).

Because the µSiM platform is a tool for modeling tissue barriers, it is necessary to establish companion protocols for assessing barrier function through transendothelial electrical resistance (TEER), small molecule permeability, or both. One challenge when working on the microscale has been adaptation of these assays for small volumes.^[53]^ The challenges have been overcome by several solutions, including incorporation of real-time measurement systems into tissue chip platforms.^[50, 54–57]^ While we have previously established methods for TEER in fully hand-built variants of the µSiM,^[25, 58]^ here we focused on developing protocols for measures of permeability by small molecule transport across monolayers. With the goal of broad dissemination of the platform, we took a fresh and more rigorous approach than our earlier examination of this topic.^[25]^ We actually developed two methods for permeability analysis via: 1) live cell microscopy (*in situ* method) and 2) conventional sampling of the ‘receiver’ chamber.^[35]^ We showed that the two methods agree both with each other and with published literature values for an immortalized brain microvascular endothelial cell line and that the sampling-based method was successfully reproduced by our collaborating UniBe laboratory.

A key innovation of the µSiM compared to all other tissue chip platforms is the use of precision-fabricated ultrathin silicon-nitride membranes. Unlike conventional thick (∼10 µm) polymer membranes, which force a choice between high porosity with poor imaging *vs.* low porosity (< 1%) with good imaging, silicon-nitride ‘nanomembranes’ have glass-like imaging despite being highly porous (∼15% porosity). This is also superior to Emulate’s commercial platform which, while an improvement from Transwell^TM^ filters, are manufactured with micron-scale (≥ 1 µm) pores and low porosities (∼1%).^[48]^ In addition, Emulate’s membranes are built entirely from PDMS, leading to concerns about small molecule adsorption.^[11, 59–60]^ In contrast, the exposed surfaces of the m-µSiM components contain only 5% of silicone material. The other materials and dimensions were selected to maintain the advantages for imaging afforded by the membranes. This includes a thin bottom component (∼0.2 mm) and the glass-like COP polymer as the bottom surface. The ability to monitor monolayer growth and maturation in live cultures without compromising membrane permeability is a significant advantage over commercial Transwell^TM^ -style products and other commercially available tissue chips, such as Mimetas OrganoPlate™^[61]^ and the SymBBB™^[62]^ products.

The ability to quickly customize the m-µSiM configuration through the exchange of components or the addition of an accessory was demonstrated in examples of: 1) side-by-side co-culture and 2) immune cell transmigration. The side-by-side culture has some advantages over the traditional “stacked” configuration, in which cell types are grown on opposing sides of the membrane.^[20, 46, 63]^ Most obvious is the ability to clearly image the two cell types as they are no longer in the same optical path, but still maintain close proximity to each other (< 1 mm). For models of the BBB, this simple reconfiguration may also be appropriate to study the interactions between the glia limitans and the BBB barriers in post-capillary venules, where astrocytic endfeet do not contact the endothelial cells.^[64]^ The similar configuration might be used to study lung-blood barrier, where an interstitial space separates barriers created by the vascular endothelial cells and lung epithelial cells.^[65–66]^ Our second example swapped our nanoporous membranes for dual-scale membranes to study immune cell migration. While quantification of migration in terms of percent migration in this study was challenging, due to the limited access of immune cells to the membrane window, this will be addressed in future studies by incorporating the flow module,^[27]^ which limits the immune cell access to just around the membrane window. We are also developing code for automated counting of migration events within videos, to improve the analytical throughput of these studies. By pairing our high-resolution videos of migration with the collection of transmigrated immune cells for down-stream analysis, the m-µSiM is uniquely positioned to test hypotheses about distinct mechanisms of migration and tissue inflammation as a consequence of immune cell migration. We can look to probe the function and consequences of cell polarity and directional inflammatory signals. With addition of other cell types, we will also have the ability test hypotheses about specific contributions of each cell type to transmigration in different disease states.

While we illustrated the use of the modular µSiM for the development of a BBB model, the platform is being actively reconfigured and expanded by multiple laboratories for a variety of forthcoming applications. Our companion paper exemplifies modular expansion of functionalities with a plug-and-play insert that provides an option to configure the core m-µSiM into a flow cell to support studies of circulating factors, including cells and physiological shear.^[27]^ This study also includes demonstration of a quick modification to Component 2 that enables use of an aligned collagen matrix in the bottom channel component for a more tissue-like substrate. In addition, unpublished studies replace Component 2 to model a healing tendon, or to reduce the working distance for very high magnifications on a µSiM nanomembrane. Importantly, the redesigned Component 2s can be rapidly prototyped, tested, mass produced, and assembled with the same workflow developed here. These and other examples lend support to our claim that the m-µSiM makes custom tissue chip design and use accessible to a broader cross-section of the biomedical research community.

## 4. Conclusion

In conclusion, we have developed and validated a modular version of our µSiM tissue chip platform to enable mass distribution, interlaboratory reproducibility, and customized design options for research laboratories interested in modeling barrier tissues. As part of the development, we have introduced two distinct methods (*in situ* and sampling) for the measurements of monolayer permeability to small molecules and demonstrated their agreement with each other. Our interlaboratory reproducibility study was done with hiPSC derived brain-like endothelial cells (EECM-BMEC protocol). Not only did the two laboratories report agreement in quantitative measurements of permeability as monolayers matured in the m-µSiM devices, they developed and executed consensus protocols for seeding, culture, staining, and imaging that will provide a foundation for future use by others. Our report also demonstrates the ability to quickly reconfigure the m-µSiM through the insertion of an accessory or the exchange of a part to enable a new tissue model or assay. Because there are limitless potential reconfigurations of the m-µSiM using existing or custom components, we intend for this feature to empower many non-engineering laboratories to design barrier models that are suited to their particular experimental needs.

## 5. Experimental Methods

### Nanomembranes

Nanomembranes are manufactured at the wafer scale (∼400 per wafer) by SiMPore, Inc. (West Henrietta, NY) and shipped from SiMPore packaged in gel boxes. In this study we use **n**ano**p**orous silicon **n**itride (NPN, SiMPore, NPSN100-1L and NPSN100-2L) membranes and dual-scale (SiMPore, NPSN100-MP-1L-3.0LP) membranes. NPN membranes are ∼100 nm thick, with ∼60 nm diameter pores and a porosity of ∼15%. Dual-scale membranes are NPN membranes with a low density of micron-sized pores to enable cell transmigration.^[18]^ The dual-scale membranes used here feature 3 µm pores that add an additional porosity of 0.625% (used by UR) or 1.25% (used by UniBe).

### µSiM Components

The top well (Component 1) and bottom channel (Component 2) of the μSiM were manufactured at ALine Inc. (Signal Hill, CA) using laser cutting and lamination processes that are compatible with mass production (hundreds to tens-of-thousands) of microfluidic devices in a single production run. The material composition is as previously described.^[22]^ While the PSA layers do contain silicone, the material accounts for only ∼5% of the fluid-exposed surface, minimizing concerns about the ability of the porous polymer to adsorb small molecules.^[59–60]^ The external surfaces of the shipped components include an additional protective layer (masking material) to maintain cleanliness and sterility during shipment and local storage. The masking material is removed by the user prior to assembly of the components. Parts were produced using a batch process and diced after final lamination for more reliable handling in the laboratory: Component 1 is shipped as single units and Component 2 is shipped as 2 x 14 strips. Individual parts of Component 2 can be removed from the strip during assembly. Component 1 contains fluidic access ports to the underside channel that create sealed fits against P20/P200 pipette tips purchased from VWR (76322-516, Radnor, PA).

In-house assembly of devices was performed in a sterile environment (i.e. biosafety cabinet). Initially, the desired membrane chip was placed on Fixture A1 (see Figure 1 for fixture definitions) in an inverted orientation using notched tweezers (758TW003, Techni-Tool, Worcester, PA). Straight-tipped tweezers (758TW534, Techni-Tool; or equivalent) were then used to remove protective masks from each side of Component 1 before the component was placed over the chip, well-side down. Fixture A2 was pressed firmly onto Fixture A1 to bond the chip to Component 1, applying pressure onto different corners of Fixture A2. Component 2 was then removed from the protective strip, and the protective layer on the opposite side was removed. Using tweezers gripping the non-adhesive corner of the component, Component 2 was placed into Fixture B1, exposed PSA and channel-side up. Component 1 was then placed onto Component 2, well-side up, and Fixture B2 was pressed firmly onto Fixture B1 to bond the two components. The assembled device was removed from the fixture, and any air bubbles on the underside of the device were pressed out using straight-tipped tweezers. Devices were further sterilized by UV for 20 minutes in the biosafety hood before using for cell culture. For more graphical/video guides to assembly see https://nanomembranes.org/resources/modular-μsim/μsim-introductory-pages/ “ALine Modular Device Assembly” and Video S1, Supporting Information.

### Cell Culture Protocols

All cell cultures were maintained in a 37°C incubator with 5% CO_2_/95% air and saturating humidity. For culture in the m-μsim, devices were kept within humidified Petri dish chambers to reduce media evaporation.

Immortalized human brain microvascular endothelial cells (hCMEC/D3) were purchased from MilliporeSigma (Burlington, MA) and used between passage 1-10 as recommended by the supplier. Cells were seeded in rat tail type I collagen (100 µg/mL, Sigma Aldrich, St. Louis, MO)-coated flasks and maintained in modified Endothelial Growth Media 2, EGM-2 (EBM-2 basal medium containing human epidermal growth factor (rhEGF), insulin-like growth factor-1 (R3-IGF-1), vascular endothelial growth factor (VEGF), human fibroblast growth factor (rhFGF-B), ascorbic acid, gentamicin sulfate/amphotericin B (GA-1000), hydrocortisone, and fetal bovine serum (FBS, 2.5%), all from Lonza Biosciences, Basel, Switzerland. Media was replaced every 2-3 days.

For functional assays with hCMEC/D3 monocultures, the top well of m-μsims was coated with a mixture of rat tail type I collagen (25 µg/cm^2^, Sigma Aldrich) and human fibronectin (5 µg/cm^2^, Corning Inc., Corning, NY) for 1 hour at 37°C, 5% CO_2_. Both chambers were then rinsed with medium, and hCMEC/D3 were then seeded on collagen/fibronectin-coated m-μsims at a density of 40,000 cells/cm^2^ in growth factor depleted EGM-2 medium, termed assay medium (EBM-2 medium containing hFGF, hydrocortisone, GA-1000, 2% FBS, all from Lonza Biosciences). Cells were allowed to settle for 2 hours, then media was replaced to remove non-adhered cells. Cells were grown 13 days, and media was replaced every 2-3 days. For more graphical/video guides to m-μsim cell culturing, see https://nanomembranes.org/resources/modular-μsim/μsim-protocols/.

IMR90-4 hiPSCs (WiCell, Madison Wisconsin) were differentiated into EECM-BMEC-like cells as previously described.^[42-43, 67-70]^ Briefly, cells were differentiated into endothelial progenitor cells (EPC), seeded at the optimized density of 100,000 cells/well in tissue culture 12 well plates on D-3. Following EPC differentiation, cells were sorted via magnetic-activated cell sorting (MACS) for CD31^+^ cells. To obtain pure EECM-BMEC-like cells, cells were selectively passaged and used in assays between passages 3-6. For assays in m-μsim, EECM-BMEC-like cells were seeded onto collagen IV (400 μg/mL, Sigma)/fibronectin (100 μg/mL, Gibco)-coated m-μSiMs at a density of 40,000 cells/cm^2^ in hECSR, which is hESFM (Gibco) with serum free B-27 Supplement (1X, Gibco) and human fibroblast growth factor 2 (20 ng/mL, R&D Systems). hECSR was added to the bottom channel prior to addition of cells into the well. Cells settled for 2 hours, then media was replaced to remove non-adhered cells. hECSR was replaced each day, and assays were performed on day 2-6 of culture. For Transwell assays, EECM-BMEC-like cells were seeded onto collagen IV (400 μg/mL)/fibronectin (100 μg/mL)-coated filters at a density of 1.12 x 10^5^ cells/filter in hECSR and grown for 6 days before measuring permeability.

Clonetics™ Normal Human Astrocytes (NHA) were purchased from Lonza Biosciences and used between passage 1-6 as recommended by the supplier. Cells were seeded in uncoated flasks and maintained in astrocyte basal medium (ABM) supplemented with rhEGF, insulin, L-Glutamine, ascorbic acid, GA-1000, and 3% FBS (all from Lonza Biosciences), termed astrocyte medium. Media was replaced every 2-3 days. For culture in m-μsim, 35,000 cell/cm^2^ were added to chambers coated with a mixture of rat tail type I collagen (25 µg/cm^2^, Sigma Aldrich) and human fibronectin (5 µg/cm^2^, R&D Systems) for 1 hour at 37°C, 5% CO_2_. Chambers were rinsed with medium prior to cell addition. Cells settled for 2 hours, then media was replaced to remove non-adhered cells. Media was replaced each day until analysis.

### In Situ Small Molecule Permeability

For cell-free assays, membranes were coated with a mixture of rat tail type I collagen (25 µg/cm^2^, Sigma) and fibronectin (5 µg/cm^2^, Corning) for 1 hour at 37°C, 5% CO_2_. For cell-seeded assays, hCMEC/D3 were seeded in the top well of coated m-μsims at 40,000 cells/cm^2^ in assay medium and cultured for 13 days. All devices were washed with fresh media prior to permeability assessment. Devices were set up on an Andor Spinning Disk Confocal microscope stage (Abingdon, United Kingdom) attached to a Nikon TiE microscope, and the detector is a Zyla 4.2 sCMOS camera. A 10X objective (NA 0.45) was focused on the nanomembrane with the membrane window in the center of the field of view along the x axis and near its center along the membrane’s long axis, and then translated down 100 µm. Media from the top well was replaced with 100 µL of the fluorescent small molecule solution (200 µg/ml 10 kDa Dextran conjugated to AF488, 1 mg/ml 10 kDa Dextran conjugated to FITC, or 150 µg/ml lucifer yellow 457 Da, all Invitrogen), and fluorescent time-lapse imaging began as immediately as possible (10 kDa Dextran conjugated to AF488: 1000 ms/image, Ex488 Em525/50BP, 5% laser power; 10 kDa Dextran conjugated to FITC: 500 ms/image, Ex488 Em525/50BP, 5% laser power; lucifer yellow: 1000 ms/image, Ex405 Em525/50BP, 35% laser power). Images were acquired once per minute for 10 minutes. Upon completion of the time-lapse imaging, media in the channel was replaced with the fluorescent solution from the top well, and a source fluorescence intensity image was acquired in the same location as the time series.

For cell-free assays, data was fit to Equation (1), a Free Diffusion equation derived from Fick’s Law, to solve for the diffusion coefficient, ***D***, and fluorescence intensity at time infinity, ***F_∞_***:

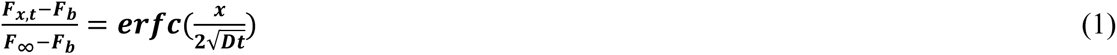

where ***F_x,t_*** is the fluorescence intensity at position ***x*** and time ***t***, ***F_b_*** is background fluorescence intensity, ***t*** is time of diffusion, and ***x*** is the distance of diffusion. For our calculations we used ***x*** = 133 µm, to account for the index mismatch caused by the refractive index change from air (1.00) to water (1.33) in the optical path.^[71]^ We experimentally confirmed that a physical shift of the objective by 100 µm moves the focal plane 133 µm.

For cell assays, data was fit to Equation (6), a Constant Flux equation, to solve for endothelial barrier permeability, ***P***. The equation was derived from the equation for molecular transport into a semi-infinite space from a boundary experiencing constant flux:^[32]^

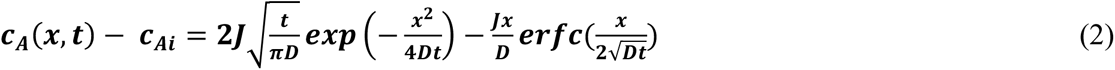

where ***c_A_(x,t)*** is the concentration of small molecule at position ***x*** and time ***t***, ***c_A,i_*** is the initial concentration at position ***x***, ***J*** is the flux of the small molecule, and ***D*** is the diffusion coefficient of the molecule. Dividing Equation (2) by ***c_0_* - *c_A,i_***, we get:

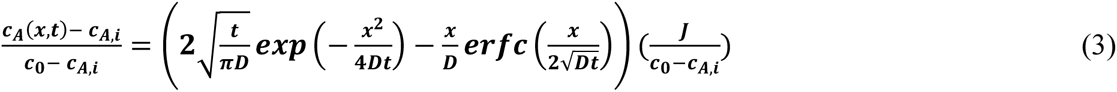

where ***c_0_*** is the source concentration of the molecule. We know that flux is equivalent to:

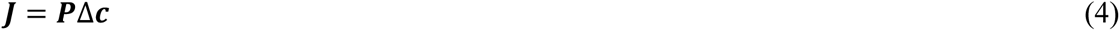

where ***P*** is permeability and ***Δc*** is difference between source concentration and the concentration of small molecule at position ***x***, time ***t***. Because ***c_0_*** >> ***c_A_(x,t)*** at time ***t*** = 10 minutes, this can be simplified to:

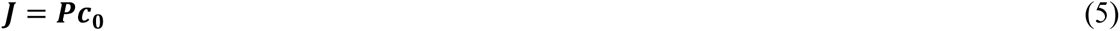

By replacement of Equation (5) into Equation (3) and assuming fluorescence is proportional to concentration, we arrive at the final equation:

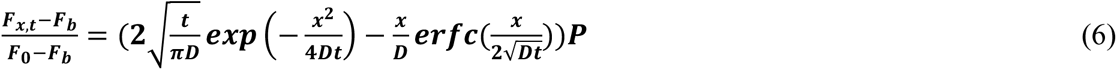

where ***F_x,t_*** is the fluorescence intensity at position ***x*** at time ***t***, ***F_b_*** is background fluorescence intensity, ***F_0_*** is source fluorescence intensity, ***t*** is time of diffusion, ***D*** is the diffusion coefficient of the small molecule, and ***x*** is the distance of diffusion. The diffusion coefficients for each molecule were calculated using the Stokes-Einstein equation at room temperature (293K) estimated with the viscosity of water at 20°C (1.003^-3^ kg·m^-1^·s^-1^) and using Stokes’ radii from the literature.^[72–73]^ Permeability, ***P***, measured through this approach represents the combination of endothelial and basement membrane permeability, but excludes the system permeability.

Data were fit to these equations using a custom-written Wolfram Mathematica code (Champaign, IL). Data for all *in situ* permeability assays were excluded if the membrane was not properly centered along its long axis in the field of view (i.e., near its top or bottom edge) or if the analytical fit exceeded a normalized root mean square error > 0.1. For cell-free fits, data was excluded when fitted values for ***F_∞_*** were >25% of a measurement made with source solutions added to the back channel at the end of the experiment.

### Sampling-Based Permeability

Sampling-based permeability assays were performed in both the m-μsim and commercial Transwells™. hCMEC/D3 or EECM-BMEC-like cells were seeded in the top well of coated m-μsims or 0.4 µm pore Transwell^TM^ filters (3401, CoStar, Washington, D.C.) in their respective medias. All assays included a coated cell-free control to determine system permeability, and fluorescence intensity was measured by a plate reader (TECAN, Männedorf, Switzerland).

For experiments using the μsim, media in the top well was replaced with 100 µL of 10 kDa Dextran conjugated to FITC (1 mg/mL, Invitrogen) or lucifer yellow, 457 Da (150 µg/mL, Invitrogen), and devices were incubated at 37°C, 5% CO_2_ for 1 hour. Following incubation, 50 µL media was added within a pipette tip attached to one port to act as a reservoir and 50 µL solution was collected by reverse pipetting from the opposite port to remove all dye from the bottom channel (see Figure 4A; Video S2, Supporting Information). System permeability, ***P_s_***, was calculated using Equation (7):^[35]^

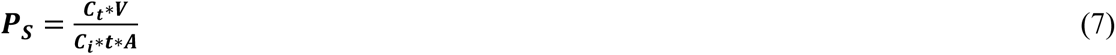

where ***C_t_*** is the concentration of fluorescent small molecule in the bottom channel at time ***t***, ***V*** is the volume transferred to the 96 well plate, ***C_i_*** is initial the concentration of fluorescent small molecule added to the top well, and ***A*** is membrane area. Endothelial permeability, ***P_e_***, was then calculated using Equation (8):

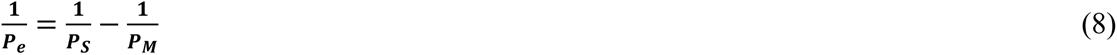

where ***P_e_*** is the permeability coefficient relating to the endothelial monolayer, ***P_S_*** is the system permeability as calculated in Equation (7), and ***P_M_*** is the system “membrane” permeability calculated on coated cell-free devices.

Commercial Transwell^TM^ clearance assays were performed as previously described.^[42]^ Briefly, media in the filter was replaced with lucifer yellow (50 µM, Invitrogen) and incubated at 37°C, 5% CO_2_ for 1 hour. Medium samples were taken from the bottom well at 15, 30, 45, and 60 minutes. An additional sample was taken from inside the filter at 60 minutes. Fluorescence intensity was then measured by a plate reader and the clearance principle was used to calculate endothelial permeability as described in detail elsewhere.^[42]^

### COMSOL Multiphysics Models

To estimate the degree of dye recovery when the backside volume is collected, we used COMSOL Multiphysics finite element software (Stockholm, Sweden) to simulate the sampling method following diffusion across a cell-free coated membrane and a cell monolayer. The two simulations involved different boundary and initial conditions as described below. Free diffusion or constant flux across the membrane was simulated for 1 hour and sample collection was modeled in both cases by introducing a 12.5 µL/s laminar inflow to one of the ports. This flow rate is based on experimental observations. For these simulations, we defined the ‘percent recovery’ as the ratio of total analytes extracted to the total transported.

#### i. Sampling after Diffusion through a Coated Membrane

We used a thin diffusion barrier boundary condition at the membrane to prevent computational load and errors generated that arise from meshing a ∼100 nm thick domain in a mm scale total volume.^[74]^ The flux across the membrane is determined using the following formula:

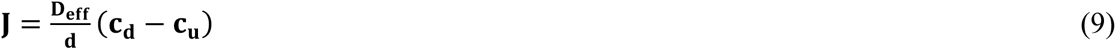

where ***J*** is flux of the analyte, ***D_eff_*** is the effective diffusion coefficient of the coated membrane, ***d*** is the thickness of the coated membrane, and ***c_d_*** and ***c_u_*** are the analyte concentrations on the two sides of the membrane. The term ***D_eff_*/*d*** can be interpreted as the mass transport coefficient, which is the reciprocal of contact resistance. The initial conditions are an initial 150 μg/mL of lucifer yellow with 100 µL in volume in the top reservoir compartment and an initial 0 µg/mL in the bottom channel compartment. The effective diffusion coefficient of lucifer yellow across the coated silicon nitride membrane was set at 1.22 × 10^-10^ m^2^·s^-1^ which was measured and calculated from previous *in situ* permeability experiments for coated membranes. The subsequent lateral diffusion in the bottom channel was simulated using Fick’s first and second laws of diffusion where the diffusion coefficient of lucifer yellow was set at 5 × 10^-10^ m^2^·s^-1^.^[75]^

#### ii. Cell-Controlled Flux

These simulations were set up in a similar fashion to (i) except we assumed the constant flux of analyte across a boundary representing the membrane area. The flux used (1.117 × 10^-7^ mol·m^-2^·s^-1^) was back-calculated from the measured endpoint concentrations of lucifer yellow in preliminary sampling experiments

### EECM-BMEC-like cell and NHA Co-culture

m-μsims were assembled using two-membrane window NPN membranes (SiMPore, NPSN100-2L). Following assembly, a cell culture insert was bonded to the chip in the open well of Component 1 using PSA (Figure 7A). To add the insert, straight-tipped tweezers were used to remove protective masks from each side of the insert and then grasped the center of the insert’s divider, PSA side down. The cell culture insert was carefully inserted into the open well of Component 1, above the membrane, aligning the center of the divider along the thick silicon between the two membrane windows. The chamber was bonded to the chip by gently pressing down with the tweezers for 2 seconds to activate the PSA. For more graphical/video guides to assembly see https://nanomembranes.org/resources/modular-μsim/μsim-protocols/ “ALine Cell Culture Chamber Insert Assembly Protocol”. The culture area of each chamber is 5.4 mm^2^ and holds a volume of ∼9 µL. Chambers were coated with the respective coating solution for each cell type and washed with hECSR. EECM-BMEC-like cells were added to one chamber at 40,000 cells/cm^2^, and NHA were seeded into the other chamber in astrocyte medium at 35,000 cells/cm^2^. Media was replaced after 2 hours. Cells were maintained for an additional 3 days in their respective mediums, with hECSR in the bottom chamber. Media was replaced each day. On day 3, excess media was added over the chambers and a glass coverslip was dropped over the device to achieve a flat imaging plane. Images were acquired on a Nikon Eclipse Ts2 phase contrast microscope.

### Immune Cell Transmigration

For neutrophil (PMN) transmigration studies at UR, EECM-BMEC-like cells were cultured in m-μsims featuring 0.625%, 3 µm dual-scale membranes using the protocols described above. PMNs were isolated as previously described.^[18, 26, 74]^ Briefly, venipuncture-derived whole blood from consenting healthy donors was deposited into sodium heparin collection tubes (BD Biosciences, Franklin Lakes, NJ) and pelleted at 500 × g for 30 minutes, 20°C with equivalent volumes of 1-Step Polymorphs solution (Accurate Chemical & Scientific Co, Westbury, NY). Following centrifugation, PMN-rich density separation layers were extracted and washed twice via centrifugation (350 × g, 10 min, 20°C). The washed PMN-rich fluid was depleted of red blood cells with an RBC lyse, followed by pelleting (350 × g, 10 min, 20°C), resuspension, and a final wash with PMN isolation buffer. Fully isolated PMNs were resuspended in 1 mL isolation buffer and left on a rotating stand to ensure minimal settling. The modular flow insert used to introduce PMNs is described in a companion paper to this one^[27]^ and enables clearer imaging of leukocytes held in a stage top incubator at 37°C compared to imaging in the open well of the m-μsim. The top flow channel was perfused with hECSR and the bottom chamber with N-Formylmethionyl-leucyl-phenylalanine (fMLP, 10 nM, Sigma Aldrich), a potent neutrophil chemoattractant. PMNs were then added into the flow chamber at 3 million PMNs/mL in hECSR via pipette injection, and transmigration was recorded on a Nikon Ti2 Eclipse inverted microscope (Nikon Corporation, Tokyo, Japan) and Zyla sCMOS (Andor Technology, Belfast, United Kingdom) using a long working distance 40X objective (NA 0.55) in phase contrast. Imaging was performed at 0.25 Hz for 30 minutes.

For T-cell migration studies at UniBe, EECM-BMEC-like cells were cultured in m-μsims on 1.25%, 3 µm dual-scale membranes using the protocols described above. CD4^+^ T helper 1 (Th1) cells were sorted and prepared for the experiment as previously described.^[76]^ Briefly, peripheral blood mononuclear cells (PBMCs) were isolated with Ficoll-Paque Plus (GE Healthcare, Chicago, IL) from human blood. Total CD4^+^ T-cells were enriched by positive selection using anti-CD4 magnetic microbeads (Miltenyi Biotech, Bergisch Gladbach, Germany). After gating on CD45RA^−^ CD8^−^ CD14^−^ CD16^−^ CD19^−^ CD25^−^ CD56^−^ cells for memory Th cells, the Th1 cell subset was sorted based on CXCR3^+^ CCR4^−^ CCR6^−^ expression and cryopreserved.^[77–78]^ Human Th1 cells were thawed 2 days prior to experimentation. Just prior to experimentation, Th1 cells were labeled for 30 minutes with CellTracker™ Green CMFDA (1 µM, Life Technologies, Carlsbad, CA). Th1 cells were then washed, live cells were collected via a Ficoll-Hypaque gradient (805 × g, 20 min, 20°C), and washed an additional two times. Th1 cells were added above the endothelial layer in 100 µL of hECSR at ∼15,000 cells/device and incubated for 2 hours. Following incubation, Th1 cells in the bottom channel were collected using the method described in the sampling permeability assay, in which a 50 µL reservoir pipet tip is added to one port and the channel solution is reserve pipetted out from the opposite port. Samples were then brought up to 150 µL and counted via flow cytometry (Figure S7, Supporting Information). Flow cytometry data was analyzed using FlowJo (Ashland, OR). Membranes with endothelial cells and adhered T-cells were then fixed with 4% paraformaldehyde (PFA) for 10 minutes and washed 3 times with PBS prior to imaging using a 10X objective (NA 0.30) on a Nikon Eclipse E600 Microscope (Nikon Corporation).

### Immunofluorescence Staining

For monoculture staining, EECM-BMEC-like cells were cultured in m-μsims in hESCR. For cytokine stimulation, media in the top compartment was replaced with recombinant human TNFα (0.1 ng/mL, R&D Systems, 210TA) and recombinant human IFNγ (2 IU/mL, R&D Systems, 285IF) for 16-20 hours. For VCAM-1 stains, stimulation was performed by replacing the media in the top compartment with smooth muscle-like cell conditioned medium (SMLC-CM) containing the same concentration of cytokines for 16-20 hours.^[43]^ For components of adherens and tight junctional complexes, cells were fixed with pre-cooled (−20°C) methanol in both the top well and bottom channel for 20 seconds and washed 3 times with PBS. The top well and the bottom channel were then blocked for 30 minutes at room temperature (RT) with 10% goat serum (Invitrogen, Waltham, MA) containing 0.1% Triton X-100 (UR and for occludin in UniBe) or 5% skimmed milk containing 0.1% Triton X-100 (UniBe). Cells were stained with primary antibodies for VE-cadherin, PECAM-1, claudin-5, occludin, and ZO-1 diluted in blocking solution for 1 hour at RT. For the live cell adhesion molecule staining, live cells were first stained with primary antibodies for ICAM-1, ICAM-2, and VCAM-1 diluted in hECSR and incubated for 15 minutes at 37°C, 5% CO_2_. Cells were then washed with PBS and the top well and channel were fixed with 4% PFA for 10 minutes at RT. Both compartments were washed 3 times with PBS, then the top well was blocked for 30 minutes at RT with 10% goat serum (UR) or 5% skimmed milk (UniBe) containing 0.1% Triton X-100. All devices were then stained with secondary antibodies diluted in blocking solution for 1 hour at RT. Nuclei were stained with Hoechst 33342 (UR) or DAPI (UniBe). Images were acquired using an Andor Spinning Disk Confocal using a long working distance 40X objective (UR, NA 0.45) or Nikon E600 Fluorescence microscope using a 10X objective (UniBe, NA 0.30) and processed using FIJI (ImageJ) software. For occludin image acquisition at UniBe, the gain function of the NIS-Elements (Basic) software was set at 19.2x. Staining was still visible in the junctions without the gain increase but was used to reduce background signal (data not shown). For UniBe images, original images were digitally cropped into quarters. For non-stimulated versus stimulated images, matching images were linearly adjusted for equivalent intensity in Adobe Photoshop (San Jose, CA) (UniBe) or FIJI (UR).

For coculture staining, EECM-BMEC and NHA were cultured as described above. Cells were fixed with pre-cooled (−20°C) methanol in all chambers for 20 seconds and washed 3 times with PBS. The top well was then blocked for 10 minutes at RT with 10% goat serum containing 0.3% Triton X-100 and stained with primary antibodies for claudin-5 and GFAP for 1 hour at RT. Top chambers were then washed 3 times with PBS and stained with secondary antibodies diluted in blocking solution for 1 hour at RT. Nuclei were stained with Hoechst 33342. Images were acquired using an Andor Spinning Disk Confocal and processed equivalently using FIJI (ImageJ) software.

A complete list of antibodies can be found in Table S2, Supporting Information.

### Statistics

For all statistical analysis, GraphPad Prism software (GraphPad, La Jolla, CA) was used. For comparison between groups with one independent variable, an ordinary one-way ANOVA was used. A two-way ANOVA was used to make comparisons for data with two independent variables, followed by a Tukey test to directly compare between groups. For the interlaboratory study of EECM-BMEC-like cell permeability, a two-way ANOVA was used, excluding UR m-μsim 6 day culture since there were no matching UniBe data. In addition, an ordinary one-way ANOVA was used to compare data within UR to include UR m-μsim 6 day culture in the analysis. P ≤ 0.05 was considered statistically significant.

## Acknowledgements

M.C.M., S.D.A., K.C., M.M, S.D., V.A, and J.L.M. were supported on NIH R61HL154249 and NIH UG3TR003281. P.K., S.H.S., B.H.S., K.K., R.E.W., and B.E. were supported on R61HL154249. S.H.S. and K.K. were partially supported on NSF CBET 1931905. H. N. was supported on the Uehara Memorial Foundation, JSPS KAKENHI Grant and the NOVARTIS Foundation (Japan) for the Promotion of Science. K.F.W. was supported by a Royal Academy of Engineering/EPSRC Postdoctoral Fellowship (EP/G058121/1) and funded by the UKRI Biotechnology and Biological Sciences Research Council (grants no. BB/K010212/1, BB/M027848/1). We thank the High Content Image Core (University of Rochester) for help with fluorescent imaging and use of the Dragonfly Spinning Disk Confocal. Microscopy at University of Bern was performed on equipment supported by the Microscopy Imaging Center (MIC), University of Bern, Switzerland.

## Ethics Approval and Consent to Participate

PMNs purified from peripheral blood at the University of Rochester were obtained from healthy subjects under an IRB approved protocol (#00004777), which includes informed consent.

