## Supporting Information for "The Modular µSiM: a Mass Produced, Rapidly Assembled, and Reconfigurable Platform for the Study of Barrier Tissue Models *In Vitro*"

**Supplemental Data**

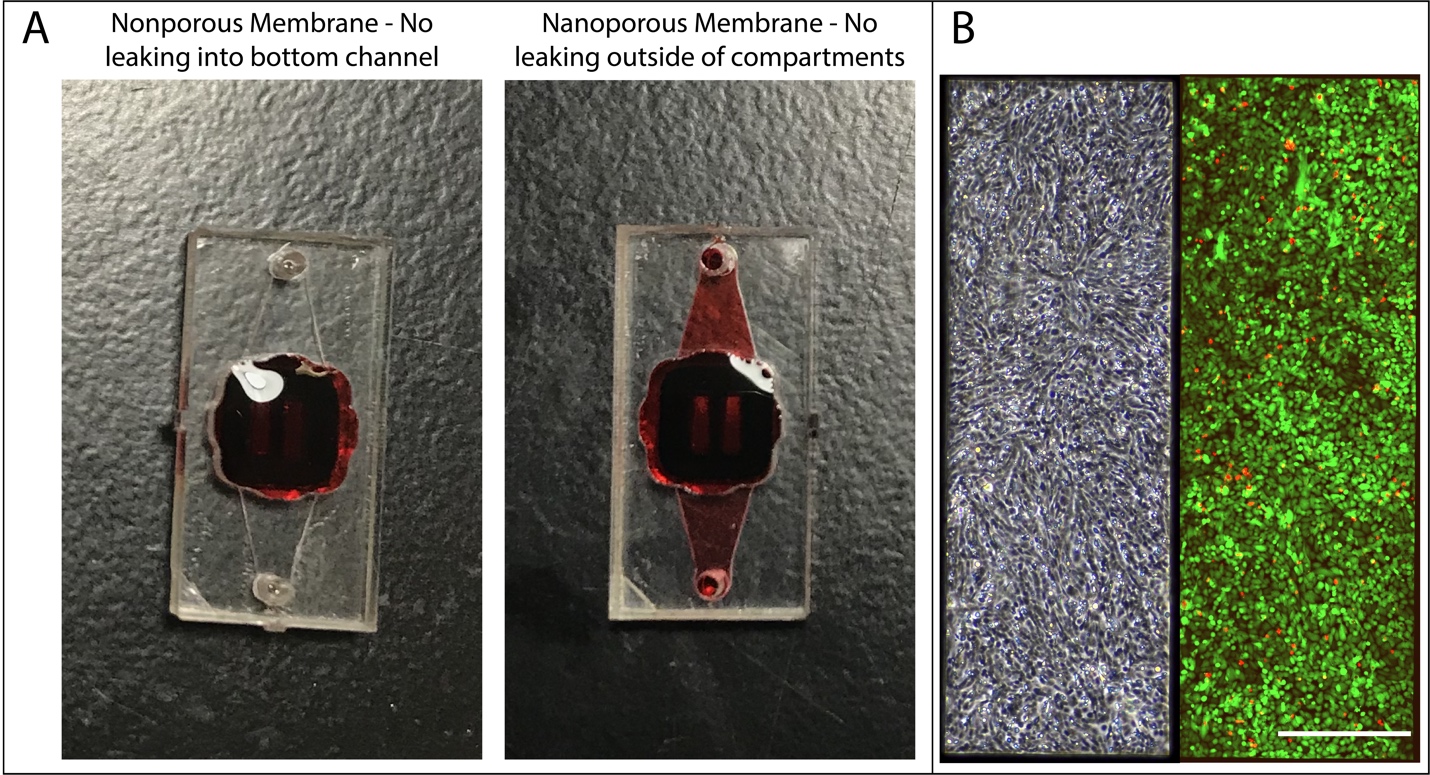
**Figure S1. m-µSiM assembly and cell culturing validation.** (A) Representative images of assembled m-μSiM devices used for a leak test. Dye was added into Component 1’s top well, and devices assembled with non-porous membranes (left) retained dye in the top well, whereas devices assembled with nanoporous membranes (right) showed diffusion into the channel after 2 hours. There was no leaking within the device. N = 4-5 devices per group. (B) Representative images of hCMEC/D3 growth in m-μSiM. Cells were cultured for 5 days. A brightfield image was taken of the monolayer (left) and a LIVE/DEAD stain was performed (right), with 98.2 ± 1.2% cell viability. Green indicates live cells, and red indicates dead cells. N = 4. Scalebar = 300 µm.

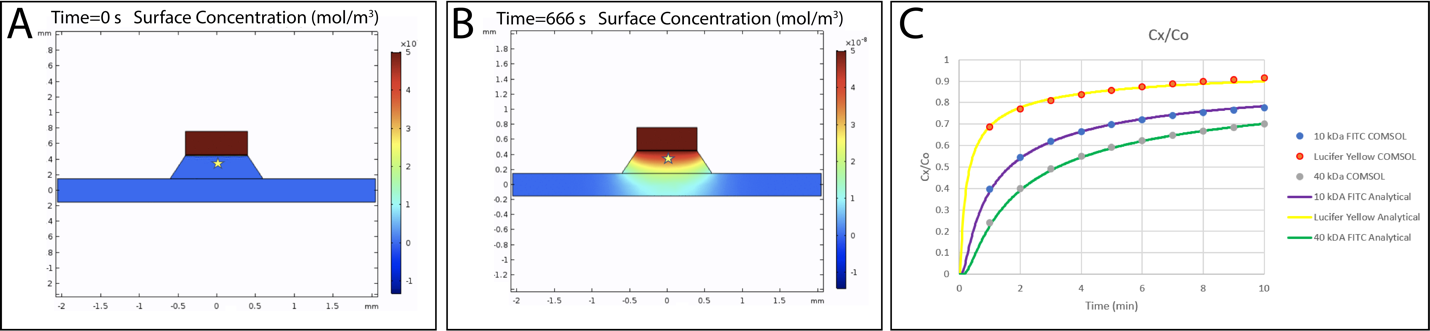

**Figure S2. Comparison between 1-D analytical model of diffusion and 2-D finite element model.** (A-B) COMSOL geometry representing membrane chip with trench, bottom chamber, and a ‘well’ of dye above the membrane at times zero (A) and 666 seconds (B). Concentration was measured at the center of the trench at a distance 100 µm below the source bottom (yellow star). (C) Concentrations over time in the center of the trench at a position of 100 µm below the membrane (C_x_) normalized to source ‘well’ concentration (C_0_). Agreement between the analytical (line) and computational (dot) models for free diffusion (Equation 1 in main text) for all molecular tracers indicates that a 1-D analytical solution can be used with the µSiM without error. Molecular tracers modeled were: 1) 10 kDa Dextran conjugated to FITC (purple line, blue dots), 2) lucifer yellow (yellow line, orange dots), and 3) 40 kDa Dextran conjugated to FITC (green line, gray dots)**.**

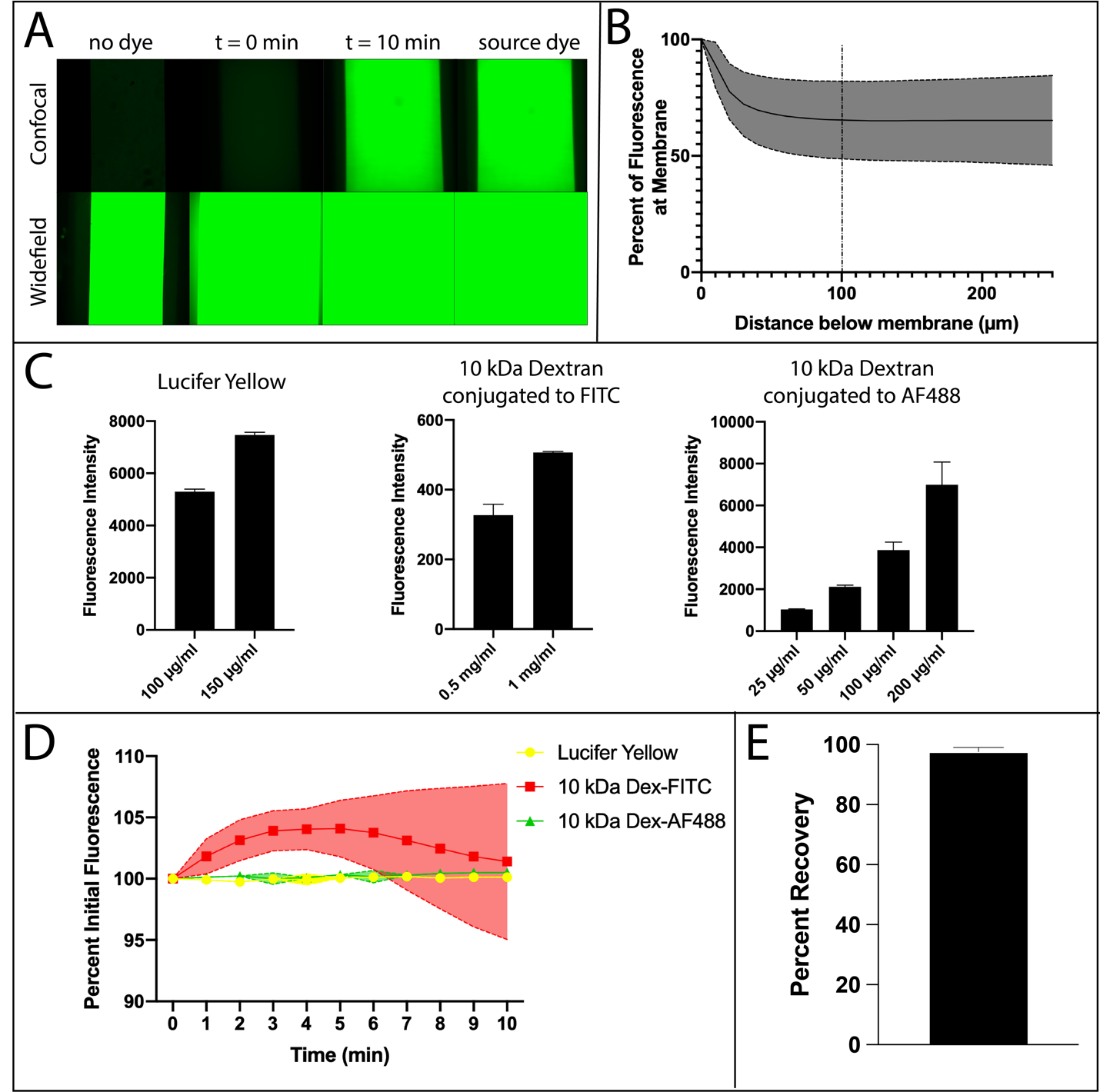
**Figure S3. *In situ* and sampling-based small molecule permeability assay optimizations.** (A) Demonstration that confocal microscopy is necessary for *in situ* permeability assay. Images are acquired on the same microscope (Andor Spinning Disc Confocal Microscope) in confocal and widefield modalities. Background fluorescence in widefield overwhelms the field of view, whereas in confocal, background is minimal and can be subtracted. (B) Optimization of confocal plane for *in situ* assay. Dye is added into the top well of a nonporous device and fluorescence intensity is measured starting at the membrane and shifting the objective down. Distance below membrane refers to distance the objective moves. At approximately 100 µm, background fluorescence from dye in the well is minimized. N = 5 devices. (C) Optimization of dye concentration for three dyes of interest in *in situ* assay. The concentration selected for each is within the linear range of fluorescence intensities to appropriately assume correlation between concentration and fluorescence. (D) Photo bleach test for three dyes of interest for *in situ* assay. 10-25% of the optimized source dye concentration is added into the channel of a nonporous device. Fluorescence intensity is measured once every minute for ten minutes. Fluorescence intensity does not diminish over the course of the experiment. (E) Experimentally tested percent recovery for sampling assay. Dye was added into the top well of uncoated nanoporous devices and allowed to diffuse for one hour. After one hour, dye was removed from the channel using a 50 µL reservoir. A second 50 µL flush was done to remove remaining dye. A fluorescent plate reader was used to measure fluorescence intensity and percent recovery was calculated as concentration in flush one divided by the sum of the concentration from both flushes. N = 10.

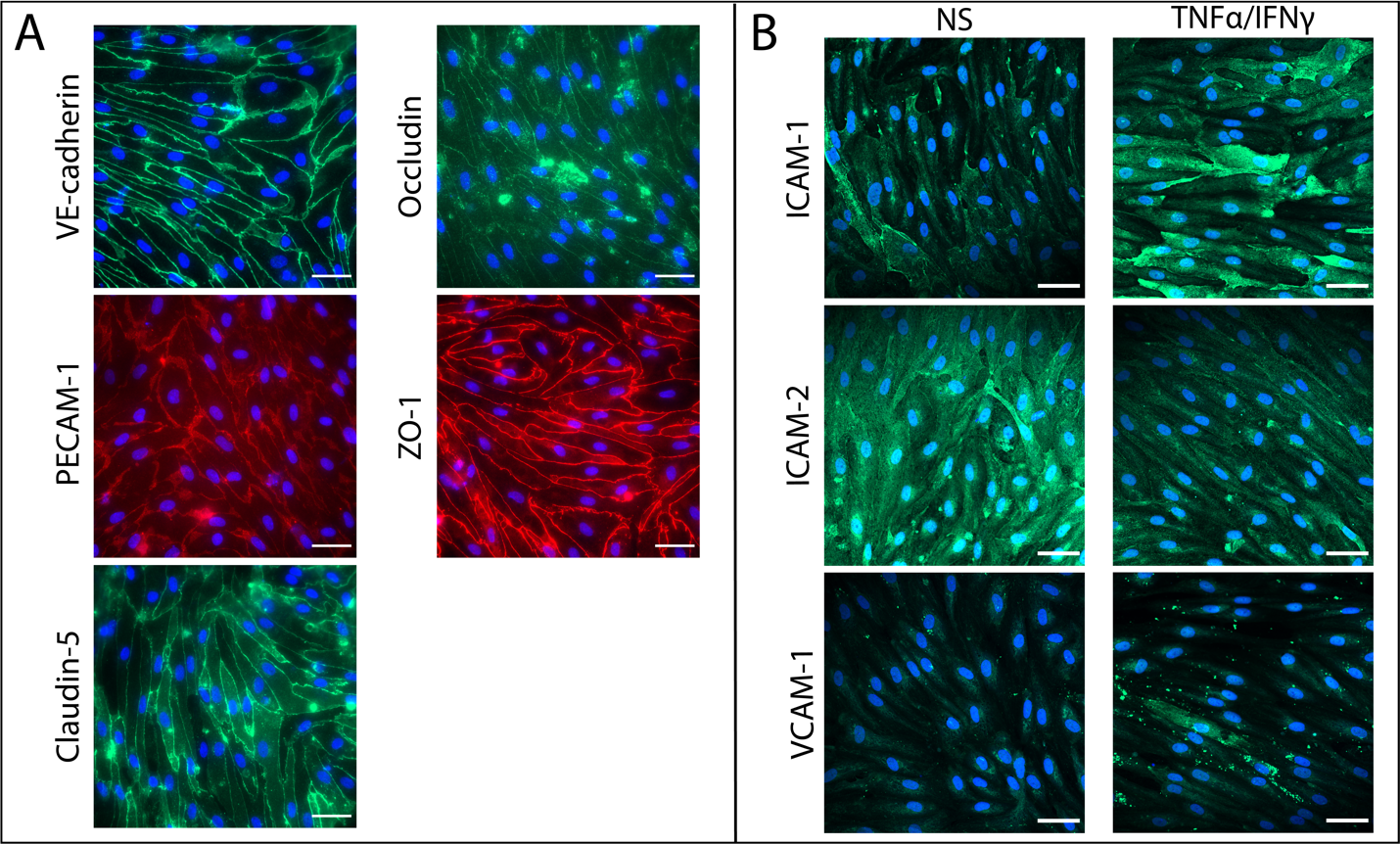

**Figure S4. Reproducibility of EECM-BMEC-like cell differentiation and culture on m-µSiM at UR.** (A) EECM-BMEC-like cells differentiated at UR and cultured on the m-µSiM express key molecules of junctional complexes, similar to those produced by UniBe (see Main Text). (B) EECM-BMEC-like cells express key cell adhesion molecules upon exposure to proinflammatory stimuli (0.1 ng/ml TNFα + 2 IU/ml IFNγ) when differentiated at UR and cultured on m-µSiM devices. Images were acquired on an Andor Spinning Disc Confocal Microscope using a long-working distance 40X objective. Scalebar = 50 µm.

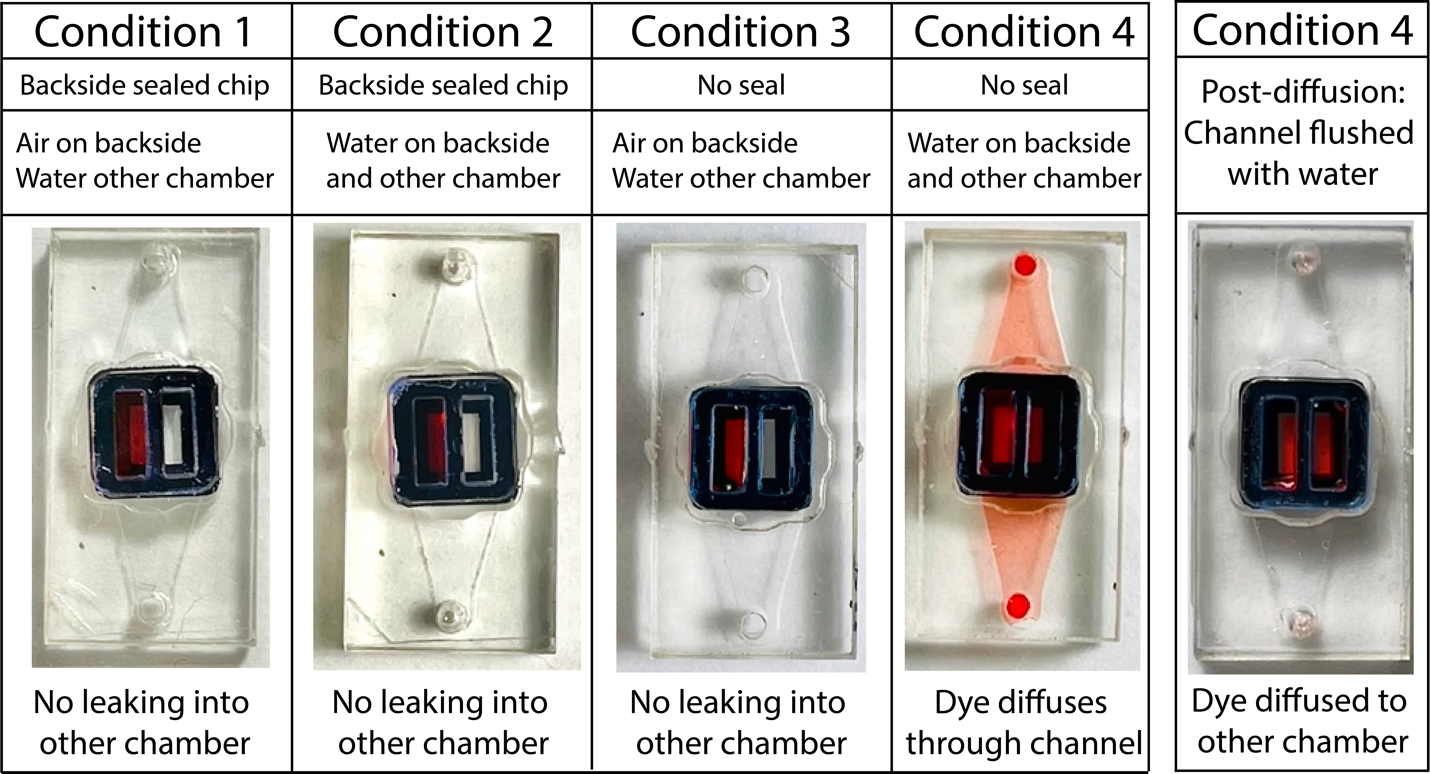

**Figure S5. Two-slot NPN insert dye leak validation.** Representative images of m-μSiM with 2-slot culture chamber insert used for a dye leak tests. Devices with the backside sealed retained dye in one chamber, with no leaking into the second chamber (Conditions 1 and 2). Devices with the backside unsealed but air in the channel (Condition 3) retained dye one chamber. When dye is added to one chamber with water in the channel and other chamber (Condition 4), dye diffuses through the channel and into the second chamber. After flushing the channel with water (Condition 4, Post-diffusion), it is clear dye does diffuse into the second chamber. This indicates communication between chambers occurs solely through the channel and not between compartments. N = 4-5 devices each group.

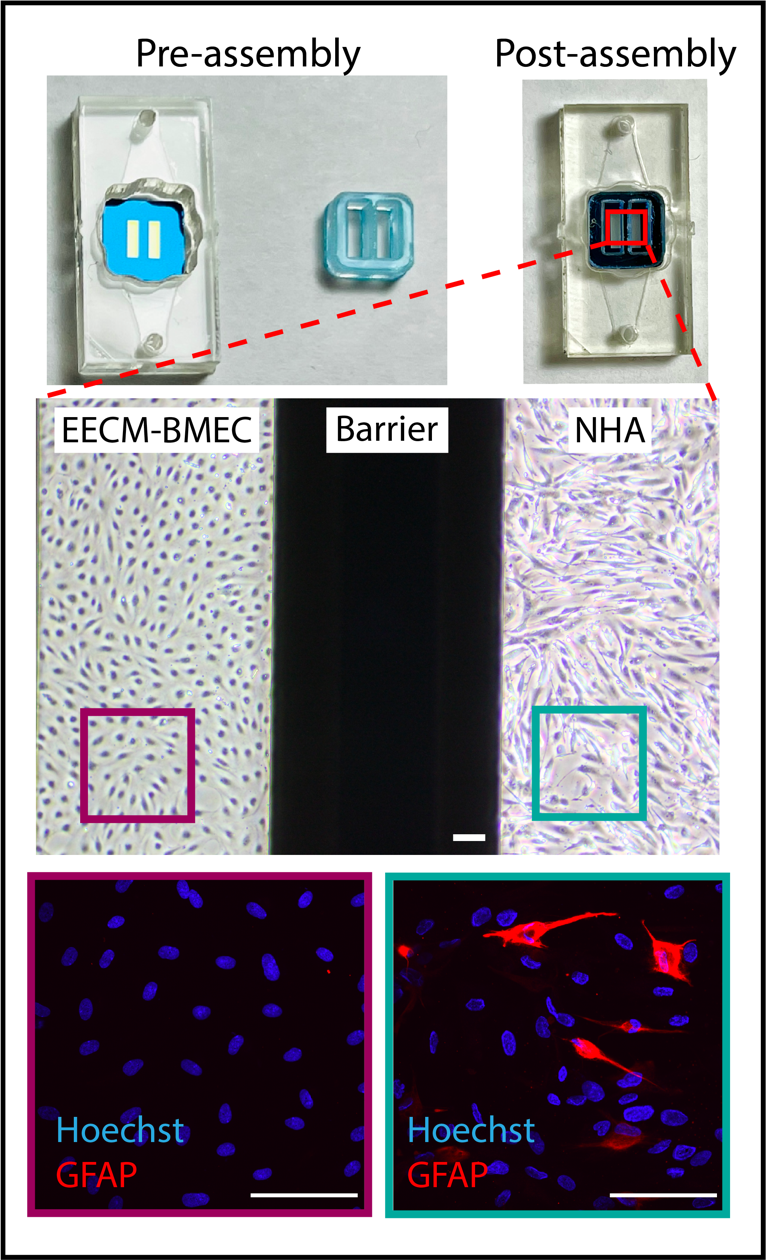

**Figure S6. Two-slot NPN insert for immunofluorescence staining of astrocytes.** EECM-BMEC-like cells were cultured in one chamber and human astrocytes (NHA) in the other chamber. Cells were stained for astrocyte marker (GFAP) and nuclear marker Hoechst (blue). Images show there is no cross-contamination of cells between culture chambers. It should be noted that GFAP is a marker of reactive astrocytes, so while it is clear through phase contrast imaging that astrocytes are growing in their respective chamber, only through staining can we see that only a handful of astrocytes are expressing GFAP in co-culture. Phase images were acquired on a Nikon Eclipse Ts2 phase contrast microscope and fluorescence images were acquired on an Andor Spinning Disc Confocal Microscope using a long-working distance 40X objective. Scalebar = 100 µm.

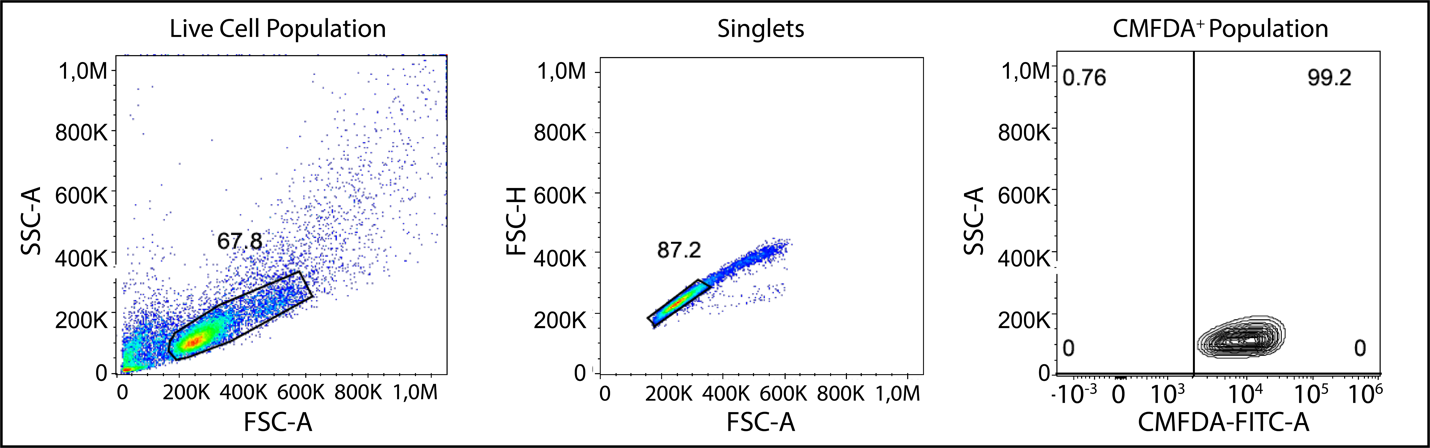

**Figure S7. Flow cytometry gating strategy for CMFDA-labelled T-cells.** Plots are initially gated to remove dead cell debris (left), then gated to remove doublets (middle). Final gate includes only CMFDA^+^ T-cells (right). Plots above are representative gating using our input T-cell population.

**Table S1. List of microscope working distances for different chip configurations.**

| **Chip Orientation** | **Distance from bottom of µSiM to Membrane**  [µm] | **Reasons to Select Orientation** | **Objectives Used in Orientation** | **Numerical Aperture (NA)** | **Working Distance (WD)**  [mm] |
| --- | --- | --- | --- | --- | --- |
| **Trench-down**  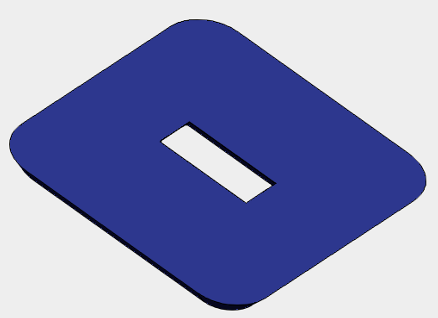  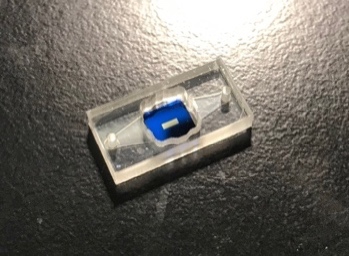 | 615 | Endothelial cells cultured in Component 1’s well grow on flat surface | 4X: Nikon, dry, Model MRH20041 | 0.13 | 16.4 |
|  |  |  | 10X: Nikon, air, Model MRD00105 | 0.45 | 4 |
|  |  |  | 10X: Nikon, air, Model MRH00105 | 0.30 | 16.0 |
|  |  |  | LWD 40X: Model, water, Part MRD77410 | 1.15 | 0.61–0.59 |
|  |  |  | LWD 40X: Nikon, air, Model MRP46402 | 0.55 | 2.1 |
| **Trench-up**  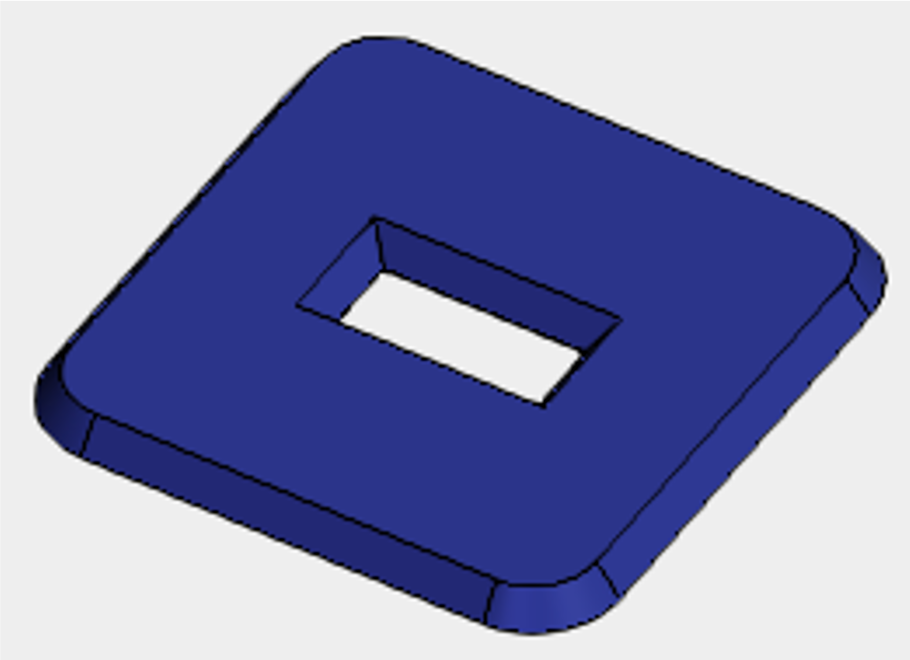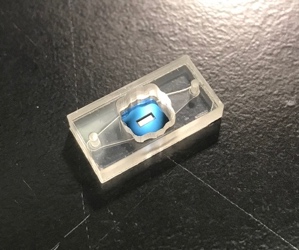 | 305 | Shorter working distance to cells and endothelial cells cultured bottom channel grow on flat surface | 20X: Nikon, multi-immersion, Model MRH07241 | 0.75 | 0.51–0.33 |
|  |  |  | LWD 40X: Nikon, water, Model MRD77410 | 1.15 | 0.61–0.59 |
|  |  |  | 60X: Nikon, water, Model MRD07602 | 1.2 | 0.31–0.28 |
|  |  |  | 60X: Olympus UPlanSApo, water, Model N1480800 | 1.2 | 0.28 |

**Table S2. Antibody list for immunofluorescence staining at UniBe and UR**

| Antibodies | Fixative | Clone | Source | Cat. N. | Secondary Antibody |
| --- | --- | --- | --- | --- | --- |
| VE-cadherin | MeOH | UniBe: F-8  UR: 123413 | UniBe: Santa Cruz  UR: R&D Systems | UniBe: sc-9989  UR: MAB9381 | UniBe: Cy3 AffiniPure F(ab')₂ Fragment Goat Anti-Mouse IgG  UR: Goat Anti-Mouse IgG (H+L) Cross-Adsorbed, Alexa Fluor 488 |
| PECAM-1 | MeOH | UniBe: MEM-05  UR: polyclonal | UniBe: Invitrogen  UR: Invitrogen | UniBe: 37-0700  UR: PA5-32321 | UniBe: Cy3 AffiniPure F(ab')₂ Fragment Goat Anti-Mouse IgG  UR: Goat Anti-Rabbit IgG (H+L) Cross-Adsorbed, Alexa Fluor 568 |
| Claudin-5 | MeOH | 4C3C2 | Invitrogen | 35-2500 | UniBe: Cy3 AffiniPure F(ab')₂ Fragment Goat Anti-Mouse IgG  UR: Goat Anti-Mouse IgG (H+L) Cross-Adsorbed, Alexa Fluor 488 |
| Occludin | MeOH | OC-3F10 | Invitrogen | 33-1500 | UniBe: Cy3 AffiniPure F(ab')₂ Fragment Goat Anti-Mouse IgG  UR: Goat Anti-Mouse IgG (H+L) Cross-Adsorbed, Alexa Fluor 488 |
| ZO-1 | MeOH | polyclonal | Invitrogen | 40-2200 | UniBe: Cy3 AffiniPure F(ab')₂ Fragment Donkey Anti-Rabbit IgG  UR: Goat Anti-Rabbit IgG (H+L) Cross-Adsorbed, Alexa Fluor 568 |
| ICAM-1 | live | HA58 | Biolegend | 353102 | UniBe: Goat Anti-Mouse IgG (H+L) Highly Cross-Adsorbed, Alexa Fluor 488  UR: Goat Anti-Mouse IgG (H+L) Cross-Adsorbed, Alexa Fluor 488 |
| ICAM-2 | live | CBR-IC2/2 | FITZGERALD | 10R-7606 | UniBe: Goat Anti-Mouse IgG (H+L) Highly Cross-Adsorbed, Alexa Fluor 488  UR: Goat Anti-Mouse IgG (H+L) Cross-Adsorbed, Alexa Fluor 488 |
| VCAM-1 | live | 51-10C9 | BD Biosciences | 555645 | UniBe: Goat Anti-Mouse IgG (H+L) Highly Cross-Adsorbed, Alexa Fluor 488  UR: Goat Anti-Mouse IgG (H+L) Cross-Adsorbed, Alexa Fluor 488 |
| GFAP | MeOH | EP672Y | abcam | ab33922 | UR: Goat Anti-Rabbit IgG (H+L) Cross-Adsorbed, Alexa Fluor 568 |

**Video S1:** Modular µSiM assembly instructions

**Video S2:** Sampling method reverse pipetting technique

**Video S3:** UR Transmigration video

<https://nanomembranes.org/the-modular-µsim-a-mass-produced-rapidly-assembled-and-reconfigurable-platform-for-the-study-of-barrier-tissue-models-in-vitro-supplemental-videos/>
